# A minimally disruptive method for measuring water potential in-planta using hydrogel nanoreporters

**DOI:** 10.1101/2020.05.29.122507

**Authors:** Piyush Jain, Weizhen Liu, Siyu Zhu, Jeff Melkonian, Duke Pauli, Susan Jean Riha, Michael A. Gore, Abraham D. Stroock

**Author notes:** Contributed equally to this work (Piyush Jain, Weizhen Liu). School of Plant Sciences, University of Arizona, Tucson, AZ 85721, USA.

## Abstract

Leaf water potential is a critical indicator of plant water status, integrating soil moisture status, plant physiology, and environmental conditions. There are few tools for measuring plant water status (water potential) in situ, presenting a critical barrier for the development of appropriate phenotyping (measurement) methods for crop development and modeling efforts aimed at understanding water transport in plants. Here, we present the development of an in situ, minimally-disruptive hydrogel nanoreporter (AquaDust) for measuring leaf water potential. The gel matrix responds to changes in water potential in its local environment by swelling; the distance between covalently linked dyes changes with the reconfiguration of the polymer, leading to changes in the emission spectrum via Fluorescence Resonance Energy Transfer (FRET). Upon infiltration into leaves, the nanoparticles localize within the apoplastic space in the mesophyll; they do not enter the cytoplast or the xylem. We characterize the physical basis for AquaDust’s response and demonstrate its function in intact maize (*Zea mays* L.) leaves as a reporter of leaf water potential. We use AquaDust to measure gradients of water potential along intact, actively transpiring leaves as a function of water status; the localized nature of the reporters allows us to define a hydraulic model that distinguishes resistances inside and outside the xylem. We also present field measurements with AquaDust through a full diurnal cycle to confirm the robustness of the technique and of our model. We conclude that AquaDust offers potential opportunities for high-throughput, field measurements and spatially resolved studies of water relations within plant tissues.

## Introduction

Plant life depends on water availability. In managing this demand, irrigated agriculture accounts for 70% of all human water use.^1^ Physiologically, the process of transpiration (*E*) dominates this demand for water (Fig. 1A): solar thermal radiation and the unsaturated relative humidity in the atmosphere drive evaporation from the wet internal surfaces of leaves; this water loss pulls water up through the plant’s vascular tissue (xylem) and out of the soil. This flow occurs along a gradient in the chemical potential of water, or water potential, *ψ* [MPa].^2^ Studies of water relations and stress physiology over the past decades have found that values of *ψ* along the path of *E* (the soil-plant-atmosphere continuum - SPAC) correlate with plant growth, crop yield and quality, susceptibility to disease, and the balance between water loss due to *E* and the uptake and assimilation of carbon dioxide (water-use efficiency).^3–5^

**Figure 1:**
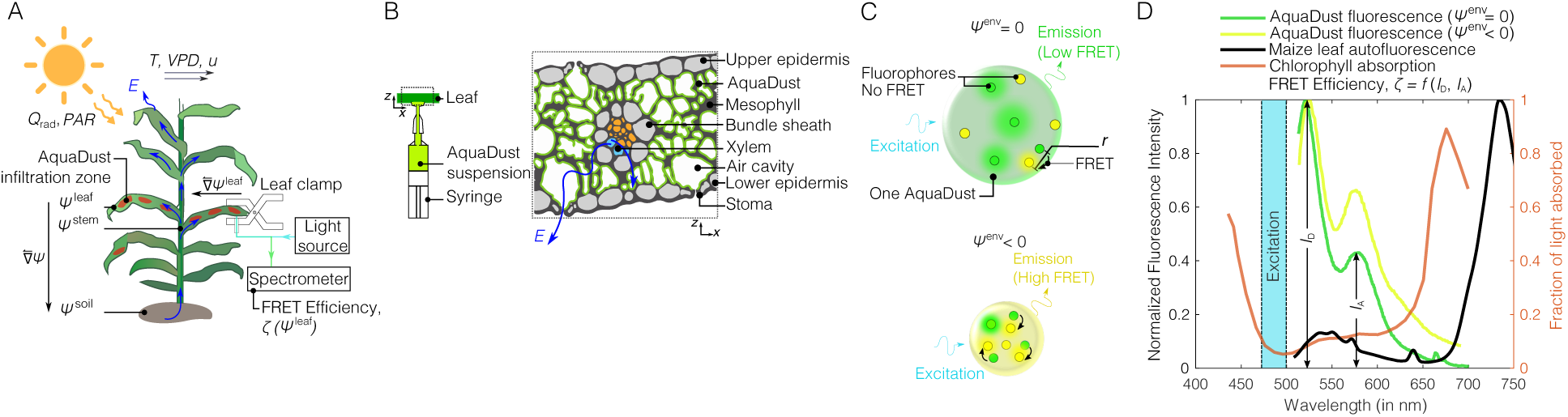
AquaDust as an in situ reporter of water potential (*ψ*): (A) Schematic representation of a maize plant undergoing transpiration (*E*) in a dynamic environment by solar thermal radiation (*Q*_rad_) and photosynthetically active radiation (*PAR*), wind speed (*u*), temperature (*T*), vapor pressure deficit (*V PD*), and soil water potential (*ψ*^soil^). Water flows through the plant (blue arrows) along a gradient in water potential 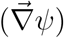. Zones on the leaves infiltrated with AquaDust serve as reporters of the local leaf water potential, *ψ*^leaf^, allowing a short (∼30 sec), minimal invasive measurement of FRET Efficiency (ζ) with a leaf clamp. (B) Schematic representations of infiltration of a suspension of AquaDust and of the distribution of AquaDust within the cross-section of a leaf. AquaDust passes through the stomata and localizes in the apoplastic spaces within the mesophyll; the particles are excluded from symplastic spaces and the vascular bundle. (C) Schematic diagrams showing mechanism of AquaDust response: the swollen, ‘wet’ state when water potential in its local environment, *ψ*^env^ = 0, (i.e. no stress condition) results in low FRET between donor (green circles) and acceptor (yellow circles) dye (top); and the shrunken, ‘dry’ state when *ψ*^env^ *<* 0 (i.e. stressed condition) results in high FRET between fluorophores, thereby, altering the emission spectra (bottom). (D) Fluorescent dyes were chosen to minimize reabsorption of AquaDust emission from chlorophyll; comparison of representative fluorescent emission from AquaDust (donor peak at 520 nm and acceptor peak at 580 nm) with the absorption spectra of chlorophyll and autofluorescence of maize leaf.

Due to the recognized importance of water potential in controlling plant function, plant scientists have spent considerable effort devising accurate and reliable methods to measure water potential of the soil, stem, and leaf.^6^ Of these, plant water potentials and particularly leaf water potential (*ψ*^leaf^) represent valuable indicators of plant water status because they integrate both environmental conditions (e.g., soil water availability and evaporative demand) and plant physiological processes (e.g., root water uptake, xylem transport, and stomatal regulation).^7,8^ To date, techniques to measure *ψ*^leaf^ remain either slow, destructive, or indirect. The current tools (e.g., Scholander pressure chamber, psychrometer, pressure probe) involve disruption of the tissue, the micro-environment, or both.^9–11^ For example, the widely-used pressure chamber requires excision of leaves or stems for the measurement of *ψ*^leaf^. Other techniques, such as stem and leaf psychrometry, require intimate contact with the tissue, and accurate and repeatable measurements are difficult to obtain.^9,12^ These limitations have hindered the study of water potential gradients along the soil-plant-atmosphere continuum and the development of high-throughput strategies to phenotype based on tissue water potential.^13^ Additionally, current methods for measuring *ψ*^leaf^ provide averages over tissues in the leaf. This characteristic makes the dissection of water relations on sub-leaf scales challenging, such that important questions remain, for example, about the partitioning of hydraulic resistances within leaves between the xylem and mesophyll.^14–16^

These outstanding challenges in the measurement of water status in-planta motivated us to develop the measurement strategy presented here, AquaDust, with the following characteristics: (1) *Non-invasive:* compatible with simple, rapid, non-invasive measurements on intact leaves. Fig. 1A presents our approach in which AquaDust reporters infiltrated into the mesophyll of the leaf provide an externally accessible optical signal that correlates with the local water potential. (2) *Localized:* allowing for access to the values of water potential at a well-defined location along the path of transpiration in the leaf tissue. Fig. 1B shows a schematic representation of AquaDust particles localized in the apoplastic volume within the mesophyll, at the end of the hydraulic path for liquid water within the plant. (3) *Sensitive and specific:* capable of resolving water potentials across the physiologically relevant range (∼ −3 *< ψ <* 0 MPa) and with minimal sensitivity to other physical (e.g., temperature) and chemical (e.g., pH) variables. Fig. 1C presents a schematic representation of an AquaDust particle formed of hydrogel, a highly tunable material that undergoes a structural response to changes in local water potential (swollen when wet; collapsed when dry). We couple the swelling behavior of the particle to an optical signal via the incorporation of fluorescence dyes (green and yellow circles in Fig. 1C) that undergo variable Forster Resonance Energy Transfer (FRET) as a function of spatial separation. Fig. 1D presents typical AquaDust spectra at high (wet - green curve) and low (dry - yellow curve) water potentials. A change in water potential leads to a change in the relative intensity of the two peaks in the AquaDust spectrum, such that the relative FRET efficiency, ζ = *f* (*I*_D_, *I*_A_), can serve as a measure of water potential. (4) *Inert:* non-disruptive of the physiological properties of the leaf (e.g., photosynthetic capacity, transpiration rate, etc.).

In this paper, we present the development, characterization, and application of AquaDust. We show that AquaDust provides a robust, reproducible response of its fluorescence spectra to changes in leaf water potential in situ and across the usual physiological range. We apply our approach to quantify the spatial gradients of water potential along individual leaves undergoing active transpiration and across a range of soil water potentials. With these measurements, we show that the localization of AquaDust in the mesophyll allows us to quantify the importance of hydraulic resistances outside the xylem. We further use AquaDust to measure the diurnal dynamics of *ψ*^leaf^ under field conditions, with repeated measurements on individual, intact leaves. These measurements demonstrate the field-readiness of our techniques and validate the leaf hydraulic model we have developed. We conclude that AquaDust offers a powerful new basis for tracking, spatially and temporally, water potential in-planta to study the mechanisms by which it couples to both biological and physical processes to define plant function.

## Results and Discussion

### AquaDust design and synthesis

We provide a detailed explanation of the design and synthesis of AquaDust in the S.I. (Sec. S1-S4). Briefly, in the selection of a specific hydrogel matrix, we used literature, theory, and experimentation to guide our design: we selected poly(acrylamide), a neutral polymer with weakly temperature-dependent swelling ^17,18^ to minimize dependence on pH, ionic strength, and temperature; and we followed Flory-Rehner theory to tune the polymer fraction (see SI, Sec. S2 A,^19–24^) with the estimate of the chemical affinity of the polymer for water (i.e., the Flory-Chi parameter, *χ*) as obtained from the swelling behavior of macroscopic gels (see SI, Sec. S3, Table S1), to match the range of the swelling transition to the physiological range of water potential (0 *> ψ >* −3 [MPa]). In the selection of specific dyes for the FRET response, we chose fluorophores for which the peaks of excitation and emission fall between the peaks of absorption of chlorophyll and can be distinguished from the peak in chlorophyll autofluorescence (Fig. 1D). We used Flory-Rehner theory and dipole-plane FRET model to iteratively find an optimal combination of monomer and cross-linker concentration, with fixed dye concentration, to maximize ζ in the range of 0 *> ψ >* −3 [MPa] (see SI - Sec. S2 A-C, Fig. S1).^25–28^ Importantly, we found that a combined theory based on Flory-Rehner swelling and dipole-plane FRET interactions allowed us to describe the calibration function, ζ(*ψ*) with a single adjustable parameter (the effective inter-dye separation in the swollen state) (see SI - Sec. S2 D, S3, and Fig. S2, S3). The robustness of this theory allows us to calibrate AquaDust at a single point (e.g., saturation), in situ.

In defining the size of AquaDust particles, the need to deliver them through the stomata and to minimize obstruction of internal cavities within the mesophyll set a micrometer-scale upper bound; the need to accommodate FRET pairs with separations ranging from 4 to 10 nm and avoid passage through the pores of cell walls set a lower bound of ∼ 10 nm (it is reported that the nanoparticles less than 10 nm in diameter can translocate through the cell wall pores^29^). To achieve size control, we synthesized hydrogel nanoparticles using inverse microemulsion polymerization with Acrylamide (AAm) as the monomer and N-aminopropyl methacrylamide (APMA) as a primary amine-bearing comonomer for reaction with donor and acceptor fluorophores conjugated via N-hydroxysuccinimidyl ester^30–33^(see SI for details on AquaDust synthesis, Sec. S4 A-B, Fig. S4). We chose an appropriate water-to-oil ratio and surfactant concentration to regulate the size of the aqueous core of the reverse micellar droplets.^34^ After synthesis, the size of these nanoparticles was 42 nm (number-averaged mean) with a standard deviation of 13 nm, as measured using the dynamic light scattering technique (see SI, Sec. S4 C, Fig. S5).

### AquaDust characterization and in situ calibration

We used maize (*Zea mays* L.) as the model species for characterization of AquaDust. Maize is one of the three most important cereal crops for world food security; knowledge of its water-stress physiology is key to improving drought tolerance.^35–37^ We infiltrated AquaDust in the maize leaves by injecting the suspension with pressure through the stomata on the abaxial surface of the intact leaf (Fig. 1B). We used AquaDust concentration of 6.6 *×* 10^8^ particles ml^−1^ with deionized water as solvent. We selected this concentration such that AquaDust fluorescent intensity was 10 fold higher than the chlorophyll autofluorescence ensuring high signal to background intensity. We used deionized water as the suspension medium to minimize particle aggregation prior to infiltration (see SI - Sec. S4 C for details).

Immediately, after the infiltration, the zone into which the suspension permeated appeared dark (Fig. 2A). In maize, this zone typically extended ∼ 6 mm laterally and ∼ 40 mm axially from the point of injection; the asymmetry of this spreading is expected given the axial connectivity of vapor spaces in the mesophyll of maize leaves.^38^ We allowed the infiltrated suspension to come to equilibrium in the leaf under standard growing conditions for 24 hours before measurement of water potential; after this equilibration, the appearance of the infiltrated zone returned to that of the surrounding, non-infiltrated tissue. At the site of infiltration, we typically observed some mechanical disturbance of the cuticle. We avoided interrogating AquaDust at this spot, as discussed below.

**Figure 2:**
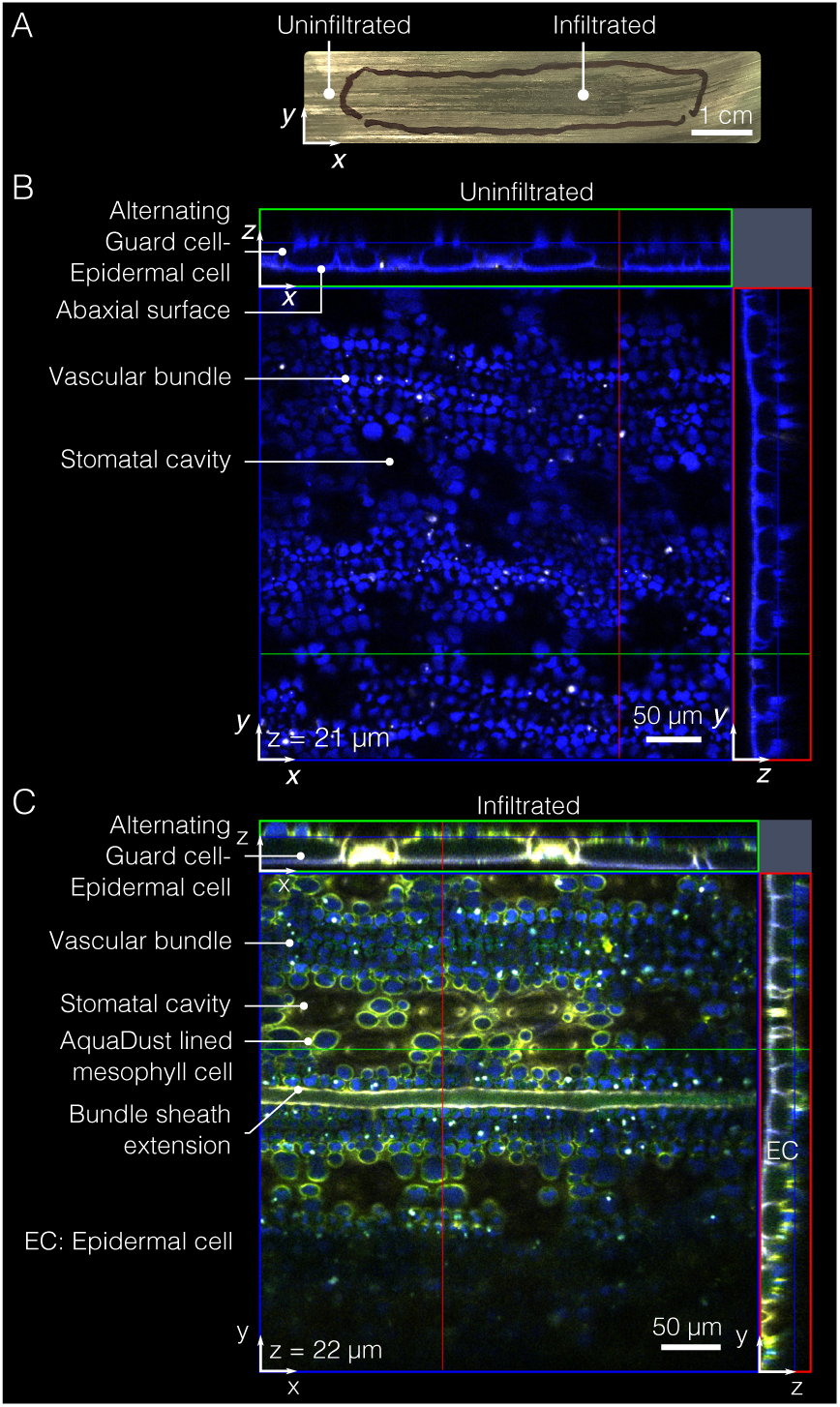
AquaDust distribution within mesophyll. (A) Typical infiltration of AquaDust suspension in maize leaf is evident with darkening of infiltrated zone immediately after infiltration; the discoloration dissipates within ∼2 hours as the injected zone re-equilibrates with the surrounding tissue (Scale bar: 1 cm). (B) Cytosol and cuticle autofluorescence (blue) from an uninfiltrated maize leaf imaged from the abaxial side using confocal microscope with xz- and yz-planes at locations denoted by green and red lines. (C) Cytosol and cuticle autofluorescence (blue) and AquaDust fluorescence (yellow) as seen from abaxial side of maize leaf under confocal microscope infiltrated with AquaDust suspension. (See SI - Sec. S4 D for details of preparation and imaging.)

Fig. 2B shows the autofluorescence from the symplasm of the bundle sheath cells and mesophyll cells (false-colored as blue), as acquired by confocal fluorescence microscopy (see SI - Sec. S4 D for details). In the top-view micrograph of the leaf without AquaDust, the autofluorescence false-colored as blue denotes the symplasm of mesophyll and bundle sheath cells^39^ (See SI - Sec. S4 D for details on sample preparation, Fig. S6 for cross-section view). In the top-view micrograph of an intact leaf infiltrated with AquaDust, the excitation of AquaDust resulted in fluorescence false-colored as yellow (Fig. 2C). We see that AquaDust co-located with the cell walls, predominantly in areas exposed to vapor pockets within the mesophyll, as seen in the micrograph shown in Fig. 2B. This distribution suggests that the AquaDust particles mostly coat rather than penetrate the cell wall. We note that we do see some evidence of penetration into non-exposed apoplastic spaces (e.g., between adjacent cells), despite the expectation that the particles ¿10 nm in diameter should be excluded from passage through cell walls.^29^ It is possible that some permeation of the nanoporous cell wall may occur due to the soft nature of the gel particles. Fig. 2C clearly shows that the AquaDust was excluded from the cytosol of all cells (mesophyll, epidermal and bundle sheath cells), and from the vascular bundles. Images of full cross-sections show this localization pattern continues through the full section of the leaf (see SI - Fig. S6 for the cross-section view of leaf with and without AquaDust). Importantly, the localization of AquaDust within the apoplast places it at the end of the transpiration path, providing an unprecedented opportunity to probe the thermodynamic state of water near the sites of gas exchange with the atmosphere.

To assess the effect of AquaDust infiltration on the physiological function of leaves, we compared the CO_2_ and water vapor exchange rates between areas of maize leaves with and without infiltration of AquaDust. We observed no significant impact of AquaDust on leaf physiological parameters (see SI, Sec. S4 E, Table S2). In order to deploy AquaDust in living plant tissues as a reporter of water potential, it is crucial to minimize AquaDust response to other physical and chemical variables such as temperature and pH. We found negligible changes in AquaDust emission spectra over a relatively broad temperature range (∼ 11 − 21°C) (see SI - Sec. S4 F, Fig. S7).^40^ This observation is consistent with the reported studies on negligible change in swelling of acrylamide gel in response to temperature.^17,18^ Also, the normalized AquaDust emission spectra were relatively insensitive (within the uncertainty range of water potential measured using AquaDust, as described in next section) over a pH range of 5-11 because of the use of non-ionic, unhydrolyzed polyacrylamide gels in the synthesis of AquaDust (see SI - Sec. S4 G, Fig. S8).^41–43^

### In-planta measurements and calibration

In order to perform non-invasive interrogations of the state of AquaDust within the leaf tissue, we developed the platform illustrated in Fig. 3A: we used an excitation source (mercury halide light source), appropriate excitation and collection filters, optical fiber probes, a leaf clamp designed to block the ambient light and to position the reflection probe (the leaf clamp did not bring the optical assembly into direct contact with the leaf), and a spectrometer to collect the fluorescence emission spectra (see details in SI, Sec. S4 H, Fig. S9). A typical measurement involved clamping the leaf for a duration of less than 30 seconds. Fig. 3B shows the emission spectra from AquaDust on intact maize (*Zea mays* L.) leaves as we subjected the potted plants to dry-down in order to progressively reduce *ψ*^leaf^ (for details, see SI - Sec. S4 I). We observed obvious, qualitative changes in the fluorescence spectrum from the leaf: the relative intensity of the acceptor dye at ∼ 580 nm rises significantly with decreasing *ψ*^leaf^ as measured using a pressure chamber (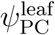, see SI- Sec. S4 I for details on how pressure chamber measurement was performed). Importantly, this large change in intensity occurs over a range of *ψ*^leaf^ typically encountered during plant water stress for most agriculturally relevant species, including maize (0 to -1.5 MPa).^44^

**Figure 3:**
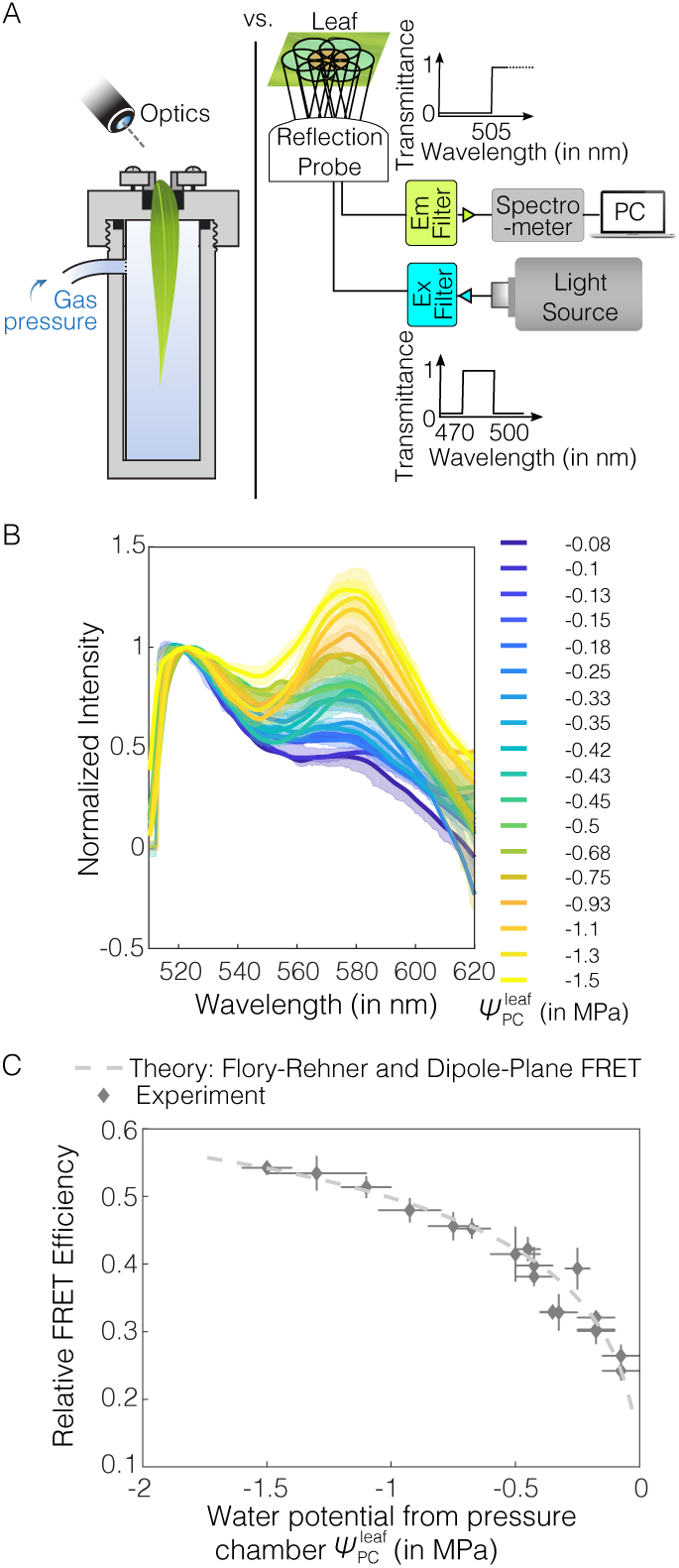
AquaDust response to leaf water potential. (A) Schematic diagrams shows calibration against Scholander pressure chamber (left) and instrumentation for a typical in situ measurement (right): A mercury lamp was used as source for illumination and a narrow-band wavelength optical filter was used to select the excitation light wavelength (here, it is 470-500 nm) used to excite AquaDust using a reflection probe. The reflected light was captured by the central fiber and sent to the spectrometer after filtering out the reflected excitation wavelengths using an emission filter to avoid the saturation of detector; spectrometer output was recorded and saved. (B) Spectra of AquaDust in maize leaves at different water potential as measured with a pressure chamber, 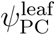 on tip of actively transpiring maize leaves. Bold lines represent spectra closest to the mean FRET Efficiency and the translucent band represents the error in the spectra as obtained from 3 to 6 measurements. The legend provides mean values of 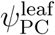 corresponding to each spectrum. (C) Relative FRET Efficiency as calculated from the spectra in (B) is plotted against 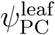. A theoretical prediction as obtained from the Flory-Rehner theory and Dipole-Plane FRET model is plotted against water potential. (see SI - Table S3 for the numerical values of the plotted data.) The vertical error bars represent range of relative FRET efficiency from AquaDust and the horizontal error bars represent range of water potential from pressure chamber.

The spectra in Fig. 3B allowed us to calibrate the AquaDust response relative to pressure chamber measurements of *ψ*^leaf^. From AquaDust emission spectra (Fig. 3B), we extracted experimental values of relative FRET efficiency, ζ_exp_, as a function of the ratio of intensity of the acceptor peak (∼ 580 nm) to that of the donor peak (∼ 520 nm) (Fig. 1D; see SI - Sec. S3 B.1). In Fig. 3C, we plot ζ_exp_ from the emission spectra (Fig. 3B) against the 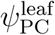(see SI - Table S3 for the numerical values). The measured values of FRET efficiencies fit a first-principles model (dashed curve) that couples the hydrogel swelling as a function of water potential (Flory-Rehner) and the FRET interaction (dipole-plane interaction;^45–50^ for details on comparison with other models,^51^ see SI - Sec. S2 B,D, Fig. S2). As with the ex situ calibration (see SI - Fig. S3), this in situ calibration involved adjusting a single parameter, *c* (separation of dyes at saturation); by fitting the theoretical FRET efficiency to the experimental FRET efficiency (ζ_th_ = ζ_exp_) at a single measurement point (here, closest to saturation: 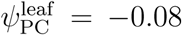 MPa), we can accurately represent the response across the full range. The requirement of a single calibration measurement limits the time required to initiate use of each new batch of AquaDust as sensor for measuring water potential. This robust, simple behavior was reproducible across the plants we have investigated (including other species such as coffee [*Coffea arabica*] and tomato [*Solanum lycopersicum* L.]) and was stable for at least 5 days in fully illuminated conditions in the greenhouse (see SI - Sec. S4 I for greenhouse conditions).

Averaged over all the readings, the difference between mean value of 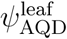, and the mean value of 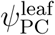, was 0.018 MPa with a standard deviation of 0.067 MPa. Based on the uncertainty associated with the experimental value of 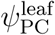 and multiple measurements from AquaDust, we found that the uncertainty in 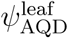 is *±*0.14 MPa based on 95% confidence interval estimate for the model compared with *±*0.05 MPa for the Scholander pressure chamber (see SI for analysis - Sec. S4 J, Fig. S10). This uncertainty is sufficiently small for most studies of water relations given the range of *ψ*^leaf^ typically encountered during plant water stress is 0 to -1.5 MPa.^52^

As noted before, we observed mechanical damage on the cuticle during injection of AquaDust by pressure infiltration (Fig. 2A); this could result in AquaDust around the site of injection being exposed to the external vapor environment. We found that the water potential reading from AquaDust was uniform and stable *±*3 mm away from the point of infiltration (see SI - Sec. S4 K, Fig. S11). As a result, measurements from AquaDust were taken ¿ 1 cm away from the site of injection to be considered as a reliable measure of *ψ*^leaf^. Since the extent of AquaDust infiltration is on the order of ≳ cm for a single infiltration in maize (Fig. 2A), the effect of damage due to injection could be reasonably avoided.

### Water potential gradients along the leaf

AquaDust opens a route to investigate local water potentials to understand and model water potential gradients in plants. As an example, we used AquaDust to track changes in *ψ* along a leaf blade to characterize key resistances to water flow in leaves.

Water moves axially from the node through the xylem and laterally from xylem into the surrounding mesophyll, down gradients in water potential resulting from the flux of water out of the surfaces of the leaf. One cause of productivity loss during drought is the increase in hydraulic resistance within the tissues that result in physiological responses (by loss of turgor, release of hormones, etc.) which can impact, for example, photosynthesis.^8,14,53^ The whole-leaf hydraulic resistance has often been measured by recording the changes in the flux of water in excised leaves with varying degrees of water stress (*ψ*^leaf^).^54^ Hydraulic decline with decreasing *ψ*^leaf^ is often attributed to the embolization of xylem vessels^55–58^ as assessed by either indirect acoustic^55,59,60^ or imaging^61–63^ techniques. However, these techniques lack quantitative information on the fraction of increase in hydraulic resistance that can be attributed to loss of xylem conductance. Recent experimental studies involving quantitative measurements of leaf xylem conductance^64,65^ and modeling studies^66,67^ have suggested that the extra-vascular resistance can contribute greater than 75% of the total leaf resistance upon dehydration. However, these experimental studies have relied on excised plant material and vein cutting (vacuum chamber method) to partition the relative roles of xylem embolism and changes in outside-xylem properties to explain the whole-leaf hydraulic decline.^64,65^ Significant uncertainty remains in the interpretation of these resistances in terms of local physiology due to the average nature of the measurement of *ψ*^leaf^ and the need to disrupt the tissue to gain hydraulic access to the xylem.^68^

Here, we used AquaDust to monitor in situ water potential gradients in an intact, mature, transpiring maize leaf during the development of soil moisture stress. Fig. 4A shows, schematically, the sites in which we infiltrated AquaDust into maize leaves for measurements of local *ψ*^leaf^ along the leaf. Fig. 4B shows the 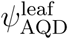 on node(*z* = *L/*6), mid(*z* = *L/*2), and tip(*z* = 5*L/*6) of the maize leaf. Under well-watered (WW) conditions, we observed a gradient ranging from 0.11 to 0.22 MPa/m from the node to tip of the leaf, with an average transpiration rate of *E* = 4.2*×*10^−5^*±*0.85*×*10^−5^ (range) kg.m^−2^.s^−1^ (see SI - Sec. S5 A for details). Similar values of transpiration-induced gradients have been reported for maize leaves, as measured using an isopiestic psychrometer (gradient of 0.17 MPa/m, *E* = 2.9 *×* 10^−5^ kg.m^−2^.s^−1^,^69^), and gradients predicted from the hydraulic architecture model for maize leaves (gradient of ∼ 0.1 MPa/m, *E* = 2.6 *×* 10^−5^ kg.m^−2^.s^−1^,^70^). Under water-limited (WL) conditions, we observed significantly larger gradients, in particular, between the mid-point and the tip of the leaves, with an average gradient of 0.7 MPa/m from the node to the tip of the leaf. This large increase in the gradient relative to the WW case suggests a substantial loss of conductance with increasing stress. Indeed, the potential drop from node to tip for a plant with limited water supply (WL) for Days 2 and 3 was 3-fold larger than that from the node to tip in a well-watered (WW) plant (Fig. 4B).

**Figure 4:**
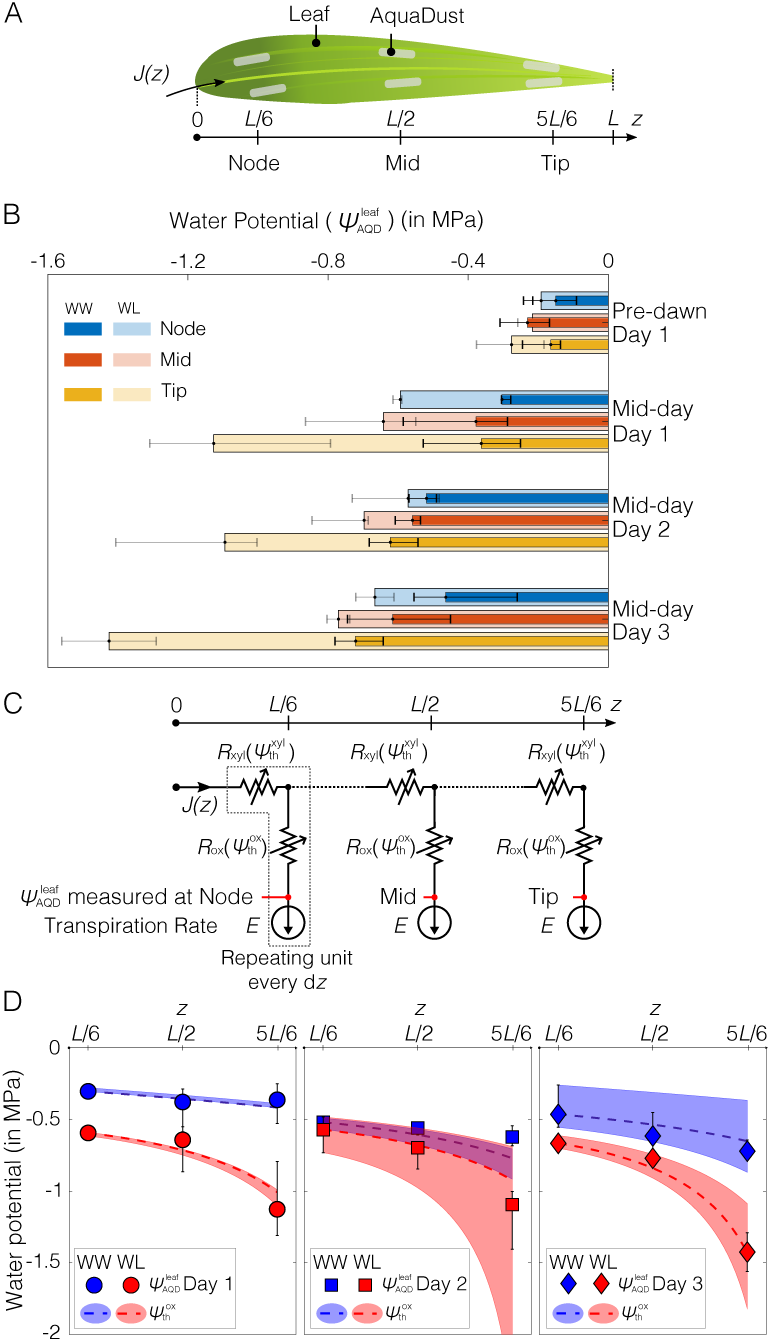
Measurements of water potential gradients along a leaf: (A) Illustration of a maize leaf with AquaDust infiltrated at the node (first one-third of leaf blade connected to stem), mid (next one-third of leaf blade) and tip (final one-third of leaf blade). (B) Water potential measured using AquaDust 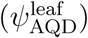 at node, mid, and tip of the leaf on maize plants in well-watered (WW) condition at pre-dawn (∼0500 hrs) and mid-day (∼1400 hrs) for three days (Day 1, 2 and 3); and for plants left unwatered (water-limited, WL) for: 1 day (Day 1) at pre-dawn (∼0500 hrs) and mid-day (∼1400 hrs), plants left unwatered for 2 days at mid-day (Day 2); and plants left unwatered for 3 days at mid-day (Day 3). Bar length and error bars represent the median and the full range respectively of water potential obtained using 3 measurements per AquaDust infiltration zone on 3 different plants. (C) Diagram of a hypothetical hydraulic circuit model of leaf with three segments (node, mid, and tip) that correspond to the sites of measurements in (B). In each segment, the resistances both in the xylem (*R*_xyl_) and outside the xylem (*R*_ox_) depend on the local xylem and outside-xylem water potential (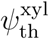 and 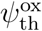). Transpiration rate (*E*) is constant and leads to a position dependent flux in the xylem, *J* (*z*). The measurements of water potential with AquaDust are assumed to correspond to 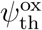 in each segment. (D) Predictions of 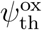 (dashed curves) with model in (C) is compared against the water potential measured using AquaDust 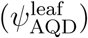 from well-watered (WW) and water-limited (WL) plants (from (B)) on three days with *E* = 4.2 *×* 10^−5^ *±* 0.85 *×* 10^−5^ kg*/*(m^2^.s); the color-coded shaded regions represent the range of values based on the range of imposed rates of transpiration. (See SI, Sec. S5 C for details of model.)

In analyzing the trends observed in Fig. 4B for WW and WL gradients, we can take advantage of the localization of AquaDust in the mesophyll, outside the xylem at the terminal end of the hydraulic pathway (Fig. 2C). This localization allows us to test hydraulic models of the intact leaf with explicit hypotheses about the partitioning of resistance between the xylem and outside xylem components of the pathway. We first tested a hypothesis in which xylem presents the limiting resistance to water flow (see SI - Sec. S5 B, Fig. S14). Starting with the magnitude and *ψ* dependence (‘vulnerability’) of xylem resistance (*R*_xyl_ (*ψ*_xyl_)) reported by Li et al.,^71^ we could not predict the variations measured with AquaDust (Fig. 4B), even with extreme adjustments of parameter values (see SI - Fig. S14). Second, we investigated the model represented in Fig. 4C in which both the resistances of the xylem (*R*_xyl_) and those outside the xylem (*R*_ox_) sit upstream of the location of our measurements with AquaDust based on the distribution we observed in Fig. 2 (also, see SI - Fig. S6).

Fig. 4C present a hypothetical hydraulic circuit model of the leaf with three segments that match our measurements at node, mid-leaf, and tip. In each segment, the xylem resistance (*R*_xyl_) and outside-xylem resistance (*R*_ox_) depend on the local values of water potential (*ψ*^xyl^ and *ψ*^ox^, respectively). We used logistic functions to represent these ‘vulnerability curves’, *R*_xyl_ *ψ*^xyl^ and *R*_ox_ (*ψ*^ox^). These logistic functions are parameterized by the WW values of resistance (*R*(*ψ* = 0)) and the potential at 50% loss of conductance (or doubling of resistance - *ψ*_50%_). For *R*_xyl_, we adopted parameter values from the literature:^70,71^ *R*_xyl_(*ψ* = 0) = 3.47 *×* 10^3^ m^2^.s.MPa*/*kg and 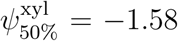 MPa. We did not find appropriate values in the literature for *R*_ox_(*ψ*_ox_) in maize.

We use this model to make predictions of *ψ*^xyl^ and *ψ*^ox^ at each segment for uniform, steady state transpiration, *E* = 4.2 *×* 10^−5^ *±* 0.85 *×* 10^−5^ kg.m^−2^.s^−1^ based on our gas exchange measurements (see SI - Sec. S5 A for details). We compared the predicted values of *ψ*_ox_ to those for measured with AquaDust, 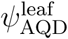. As described in detail in SI (Sec S5 C and Fig. S15), we optimized the parameters in *R*_ox_(*ψ*_ox_) to fit our measurements across both WW and WL conditions and all days. We find that this model (Fig. 4C) is consistent within the uncertainty in transpiration rate (shaded regions in Fig. 4D) with all measurements of local stress. Further, our optimal parameter values of *ψ*-dependence for extra-vascular resistance are in the range reported for the mesophyll resistance obtained for different species based on the vacuum pressure method and modeling studies.^64,65^

The agreement between 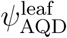 and 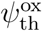 supports existing assessments of leaf hydraulics with respect to the dominance of extra-vascular resistance. The agreement between 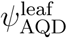 and 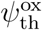 also reinforces our interpretation, based on the localization of AquaDust (Fig. 2C), that it measures outside-xylem water potential. Our observations demonstrate the capability of AquaDust to serve as an in situ reporter of local *ψ* and to help better understand the partitioning and responsiveness of resistances in leaves.

### Documentation of diurnal variation in leaf water potential in intact plants in the field

The relative rates of water loss (transpiration) and water uptake controls the water status of a plant. Evaporative demand varies with net radiation, relative humidity, air temperature and wind speed, and soil water status, as well as physiological responses of the plant resulting in fluctuations in *ψ*^leaf^ (Fig. 1A). To date, access to the dynamics of plant water stress in the field has required destructive sampling of tissues (e.g., one leaf per measurement with pressure chamber) or inference from measurements in the soil and atmosphere (eddy covariance, etc.). It is also worth noting that inference on water status from the eddy covariance method is complex and modeling requires years of effort in calibrating transpiration and canopy conductance with respect to plant water status. One of the advantages of AquaDust is that it provides minimally invasive measurements of intact plant tissues and hence, can be used for repeated measurements of water status on individual leaves to track dynamics. The response time of the AquaDust to a step change in water potential occurs on the order of seconds (See SI - Sec. S4 L, Fig. S12). The response time of leaves to the changes in environmental conditions is expected to be on the order of 15 minutes;^72^ hence, AquaDust opens opportunities to study water stress response of leaves to changing external environmental conditions. Here, we used AquaDust to measure the diurnal variation in leaf water potential and compared the predicted leaf water potential based on soil-plant-atmosphere hydraulic resistance model informed by the model in Figs. 4C-D with the measured 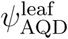 over the course of a day in field conditions.

We found general agreement between calibration of AquaDust in growth chamber and the calibration of AquaDust in field conditions (see SI - Sec. S4 M, Fig. S13 for details). Once calibrated, we documented changes in 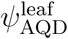 of maize leaves over a period of 15 hours in a well-irrigated field (minimal effect of soil moisture status). We performed measurements on two adjacent maize plants in an instrumented research plot at Cornell’s Musgrave Research Farm (Location: 42°43′N, 76°39′W). With AquaDust infiltration in leaves 4 and 7, we acquired three measurements per leaf once or more per hour throughout the day (except during field irrigation between 0800 and 1100 EST, Fig. 5B). We compared 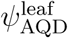 with the prediction of 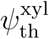 and 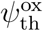 obtained using a hydraulic resistance model with the resistance from a maize leaf as shown in Fig. 4C (see SI - Sec. S5 D, Fig. S16 for details^73–75^) with the following inputs: (1) We used eddy covariance to estimate rates of transpiration (*E*; Fig. 5A); (2) we used the values of xylem resistance 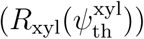and outside-xylem resistance 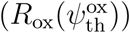 inferred from the observed gradient of water potential along the leaf in the previous section (Sec. D, Fig. 4); and, (3) we assumed that soil was saturated (*ψ*^soil^ = 0) and root and stem presented negligible resistance to water uptake under well-watered field conditions. The measurements of the diurnal dynamics of 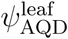 agreed favorably with the predictions of the model, further validating the model for the maize leaf (Fig. 4C) and limiting resistance for water loss being located in tissue outside the xylem. This demonstrates the potential for AquaDust to track plant water status under variable climate conditions with minimally perturbative, and allow for rapid and repeated measures of *ψ*^leaf^, and aid in more realistic modeling aimed at understanding local scale water transport in leaves.

**Figure 5:**
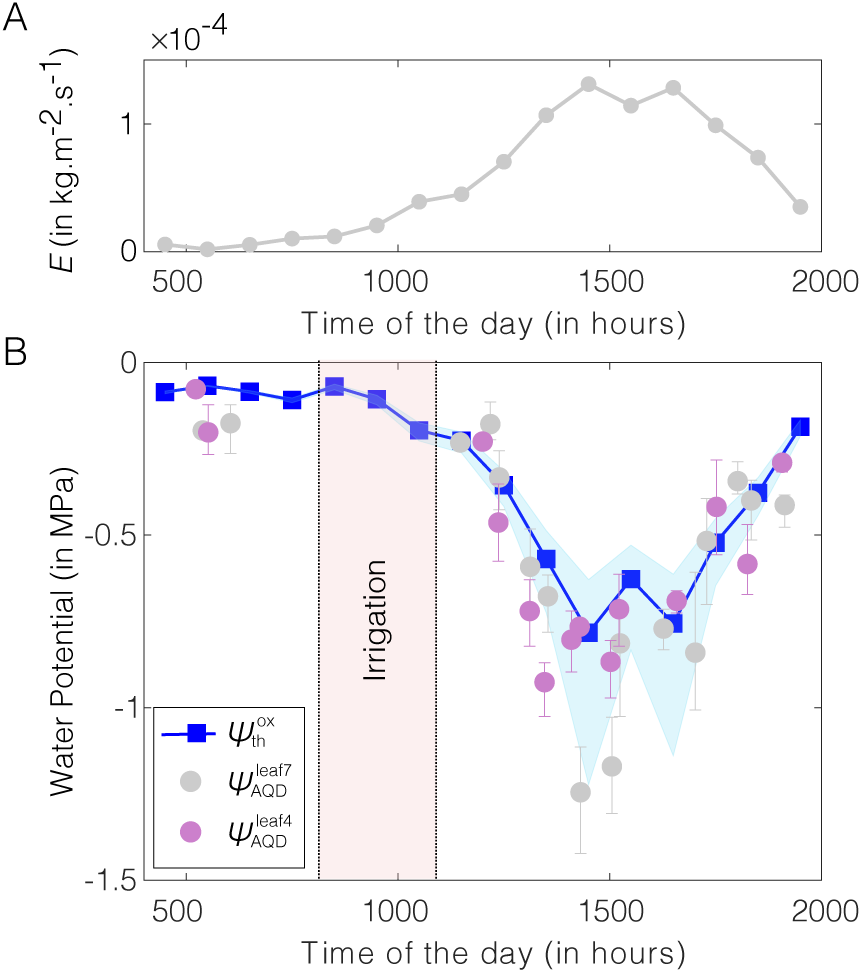
In-field diurnal measurements of leaf water potential using AquaDust: (A) Hourly-averaged transpiration (*E*) measured using eddy covariance method. (B) Values of water potential at tips of leaves 4 and leaves 7 measured with AquaDust (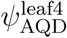, 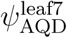) compared with the predicted diurnal variation of outside-xylem water potential 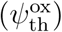 obtained using soil-plant-atmosphere hydraulic resistance model defined based on model and data in Fig. 4C-D (see SI, Sec. S5 D, Fig. S16 for details on model). Error bars represent range of water potential from two biological replicates (plants) with three measurements per replicate. Shaded blue region represents the range on theoretical prediction of 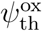 corresponding to the minimum and maximum value of outside-xylem resistance inferred from water potential gradients (shown in Fig. 4C, see SI, Fig. S15 for the numerical values of resistances).

## Conclusion

Our approach, based on hydrogel based nanosensors, AquaDust, allows for in situ, minimally invasive measurements of water potential in local physiologically-relevant micro-environments. This tool opens opportunities for better understanding of physics and biology of water dynamics in plants. As the process of AquaDust infiltration in leaves and fluorescence readout matures, AquaDust could be used for a high-throughput phenotyping strategy that allows for the discovery and quantification of new traits impacting water-use efficiency in crops. AquaDust, given its scale and localization within the mesophyll, provides opportunities to map gradients of water potential driving water flux from xylem to mesophyll and to atmosphere, and to identify the major resistances along the pathway from node to the sites of evaporation. It also opens up possibilities to address key questions that center on providing an independent estimate of the water potential of the evaporative surfaces during transpiration, critical in measurements of exchange of carbon dioxide and water vapor.^76,77^ As a tool for optical mapping of water potential, AquaDust has the potential to serve in a variety of contexts beyond leaves: in the rhizosphere, the critical root-associated volumes of soil in which water dynamics remains poorly characterized;^78^ in biophysical studies across species in which responses to local water availability are of interest;^79^ and in non-biological contexts - food science, geo-technical engineering, material synthesis - in which the thermodynamics and transport of water are important.^80–82^

## Supporting Information Available

Materials and methods for synthesis, characterization, calibration and usage of AquaDust is described in SI.

## Acknowledgement

The authors would like to acknowledge the technical assistance of Glenn Swan and Nicholas S. Kaczmar; Prof. William Philpot for providing ST2000 spectrometer and optical fiber probes; Prof. Ying Sun, Dr. Christine Yao-Yun Chang and Jiaming Wen for their technical assistance with using GFS3000 gas exchange measurement device and pressure chamber; Prof. Jocelyn K. C. Rose and Dr. Iben Sorensen for their technical support for sample preparation for the confocal imaging; Prof. Warren R. Zipfel and Dr. Rebecca M. Williams for their assistance with u880 confocal microscope imaging; Dr. Olivier Vincent for assistance in using vacuum setup; and Jacob L. Wszolek in Cornell Guterman Lab for maintaining the plants in greenhouse and growth chamber. This work was supported by the Agriculture and Food Research Initiative Competitive Grant no. 2017-67007-25950 from the USDA National Institute of Food and Agriculture, the Air Force Office of Scientific Research Grant no. FA9550-18-1-0345, and Cooperative Research Program for Agriculture Science and Technology Development (Project No. PJ01321305) Rural Development Administration, Republic of Korea. This work was performed in part at the Cornell University Biotechnology Resource Center (NIH S10RR025502 for data collected on the Zeiss LSM 710 Confocal, NIH S10OD018516 for data collected on the upright Zeiss LSM880 confocal microscope (u880)), and in part at the Cornell NanoScale Facility, a member of the National Nanotechnology Infrastructure Network (National Science Foundation, grant no. ECCS-1542081).

## Notes

### Competing Interest Statement

The authors have declared no competing interest.

