## Supplementary Information for "A minimally disruptive method for measuring water potential in-planta using hydrogel nanoreporters"

Corresponding Author: Abraham D. Stroock

### This PDF file includes:

### S1. Materials

Acrylamide (AAm) (40%($w/v$)), N,N-methylene bisacrylamide (BisAAm, $>$ 98%), Ammonium Persulfate (APS, $>$ 99.99%), Dichlorodimethylsilane (DCMS, $>$ 99%) were purchased from Sigma-Aldrich; Tetramethylethylenediamine (TEMED, Electrophoresis grade) was purchased from Fisher Scientific; N-aminopropyl methacrylamide (APMA, $>$ 98%) was purchased from Polysciences Inc; Dioctyl Sulfoccinate Sodium salt (AOT, 96%) and Polyoxyethylene(4)lauryl ether(Brij30) were purchased from ACROS Organics; n-Hexane (95%, HPLC Grade) was purchased from Millipore Sigma; N,N-Dimethyl formamide (DMF, Anhydrous) was purchased from Mallinckrodt Inc.; FRET pairs: (i) Alexa Fluor 488 (AF488) and Alexa Fluor 568 (AF568) (used as FRET pair in synthesis of bulk gels (Sec. S3)) and (ii) Oregon Green 488 N-hydroxysuccinimidyl ester (OG) and N-hydroxy succinimidyl ester Rhodamine (RH) (used in synthesis of AquaDust (Sec. S4)), were purchased from Thermo Fisher Scientific; Ethanol (Anhydrous, 100%) and Isopropyl alcohol (IPA) (99%) were purchased from VWR International; Phosphate-buffered saline (PBS) 1x tablet (10 mM Phosphate buffer, 137 mM Sodium Chloride and 2.7 mM Potassium Chloride) was purchased from Amresco; and Low-melting Agarose (LMA, Grade-Biotech) was purchased from Neta Scientific.

### S2. Theory for AquaDust response:

**A. Gel swelling behavior.** We adopt the following picture of the gel's interaction with water (1, 2): Soft polymeric matrices host solvent as a confined solution in which both molecular scale mixing and mechanical stress contribute to the energetics of solvation. The Flory-Rehner theory provides a useful, semi-empirical expression that captures these contributions (3, 4). We use modified Flory-Rehner theory proposed by Tanaka (2) to describe the free-energy of water molecules inside the gel using three terms on the right side of the following equation:

$$\psi = \Delta\mu/v_{\mathrm{w}} = \frac{RT}{v_{\mathrm{w}}}\ln(a_{\mathrm{w}}) = \\ \frac{RT}{v_{\mathrm{w}}}\left\{(\ln(1-\phi)+\phi)+\left(\frac{\theta}{2T}\phi^2\right)+\left(v_{\mathrm{w}}n_{\mathrm{o}}\left[\left(\frac{\phi}{\phi_{\mathrm{o}}}\right)^{1/3}-\frac{\phi}{2\phi_{\mathrm{o}}}\right]\right)\right\}, \quad \text{[S1]}$$

where R = 8.314 [J/K] is the universal gas constant, $T$ [K] is the temperature, $a_{\mathrm{w}}$ is the imposed vapor activity, $\phi$ is the volume fraction of monomer in final state, $\phi_{\mathrm{o}}$ is the volume fraction of monomer in the polymerization solution, $\theta$ is the flory temperature that accounts for interaction energy between polymer and solvent (Flory-Chi parameter, $\chi = \theta/(2T)$), $v_{\mathrm{w}}$ is the molar volume of solvent, (in our case, water) and $n_{\mathrm{o}}$[m$^3$/mol] is the moles of constituent chains per unit volume, $\Delta\mu$ [J] is the free energy of water molecules in surroundings relative to the reference state. In Eq. (S1), we use the approximation that the water potential of the liquid, $\psi$ [Pa] is equal to the deviation in chemical potential from saturation, $\Delta\mu$ divided by the molar volume, $v_{\mathrm{w}}$ [m$^3$/mol], or, it can be expressed in terms of the activity of the water, $a_w$ at given temperature $T$ [K] as shown in the middle expression in Eq. (S1).

*Assumptions:* For a given gel with known monomer and cross-linker concentration, we calculate the change in ratio of volume of gel ($= V/V_o \equiv \phi_o/\phi$) where $V$ and $V_o$ are volume of gel in final state and initial state during gelation respectively, where, in the initial state, the volume fraction, $\phi_o = (v_{mon1} \times w_{mon1}) + (v_{mon2} \times w_{mon2})$ (see Table S1). To calculate $n_o$, we assume that all the monomer and crosslinker introduced in the polymerization solution contribute in the formation of an homogeneous uniform hydrogel such that $n_o = \left(\frac{w_{mon1}}{mw_{mon1}} + \frac{w_{mon2}}{mw_{mon2}}\right) / \frac{w_{cl}}{mw_{cl}}$ (see Table S1 for gel composition).

**B. Models for multi-fluorophore FRET interaction.** To design an optical reporter of $\psi^{env}$, we can couple to the changes in molecular conformation implicated by changes in $\phi$: typical distances between polymeric chains vary as $\phi^{-1/3}$ for isotropic swelling of a gel. In the synthesis of AquaDust, we covalently linked a Forster Resonance Energy Transfer (FRET) pair (a donor dye and an acceptor dye) to the polyacrylamide network (Fig. 1C - Main text). Forster Resonance Energy Transfer (FRET) between pairs of fluorescent dyes can provide a spectral signature of such changes in molecular separation. Specifically, the experimental relative FRET Efficiency, $\zeta_{exp}$ is a function of the relative intensity of the fluorescence emission by the acceptor dye and the donor dye.

By design, AquaDust has multiple donors interacting in nonradiative energy transfer with multiple acceptors randomly distributed in the local environment. The dependence for energy transfer on distance between uniformly distributed multiple fluorophores in cell-membrane based FRET studies is modeled using various geometric arrangements of donors and acceptors. Here, we evaluate the predictions of different models for FRET interaction between donors and acceptors. We compare the models (Figure S2) for the multi-fluorophore interaction to the evolution of experimental FRET efficiency from AquaDust as a function of water potential.

***B.1. Dipole-dipole FRET model.*** A single donor-acceptor FRET interaction is modeled as a single pair of interacting dipoles (5):

$$\begin{aligned}\zeta_{th}(\text{Forster}) &= \frac{\left(\frac{R_o}{r}\right)^6}{1 + \left(\frac{R_o}{r}\right)^6} \\ &= \frac{\left(\frac{R_o}{r_i} \times \frac{r_i}{r}\right)^6}{1 + \left(\frac{R_o}{r_i} \times \frac{r_i}{r}\right)^6} \\ &= \frac{c'}{\left(\frac{r}{r_i}\right)^6 + c'}\end{aligned} \qquad \text{[S2]}$$

where $c' = (R_o/r_i)^6$, and $r_i$ is the initial distance between fluorophores and $R_o$[nm] is the Förster distance Typical values for $R_o$ range from $6 - 10$ nm (5), in particular, here, (i) for the FRET pair used in synthesis of macroscopic gels (Sec. S3, AF488 and AF568), $R_o = 6.2$ nm (6); (ii) for the FRET pair used in the synthesis of AquaDust (Sec. S4, OG and RH), $R_o = 5.8$ nm (7). Because of the normalization with respect to $r_i$, we do not use value for $R_o$ explicitly for calculations. By definition, $c'$ is only a function of $r_i$; experimentally $r_i$ should only depend on the concentration of dye molecules conjugated to the polymer matrix.

Assuming isotropic strain, we can express the change in distance between the dyes as function of initial and final volume of gel $(V_o, V)$ as well as the initial and final volume fraction of polymer $(\phi_o, \phi)$ as follows:

$$\frac{\phi_o}{\phi} = \frac{V}{V_o} = \left(\frac{r}{r_i}\right)^3 \tag{S3}$$

Upon substituting Eq. (S3) in Eq. (S2), we obtain

$$\zeta_{th}(\text{Forster}) = \frac{c'}{\left(\frac{\phi_o}{\phi}\right)^2 + c'}. \tag{S4}$$

Combining the Flory-Rehner model (Eq. (S1)) relation between water potential $(\psi)$ and volume fraction of polymer $(\phi)$ with Eq. (S4), we obtain a one parameter model that relates $\zeta_{th}$ with $\psi$:

$$\zeta_{th}(\text{Forster}) = f(\psi, c'). \tag{S5}$$

The parameter $c'$ is calculated by equating $\zeta_{th}$(Forster) with the experimental FRET Efficiency, $\zeta_{exp}$ closest to saturation $(\psi_{exp} = 0)$. We plot Eq. (S5) in Figure S2 for comparison with other models.

***B.2. Dipole-plane FRET model.*** As described in the main text, we have chosen dipole-plane FRET model to describe the interaction between uniformly dispersed donor and acceptor fluorophores. This model has been proposed in context of bio-membrane studies, for example, a donor-containing protein approaching acceptors embedded in an associated phospholipid mono/bi-layer(5, 8–11). The non-dimensional rate of energy transfer, $k$, between a donor separated by distance $r$[nm] from a plane of acceptors is given as:

$$k = \frac{\pi}{2} N_A R_o^6 r^{-4} \tag{S6}$$

where $N_A$[nm$^{-2}$] is the area concentration of acceptors and $R_o$[nm] is the Förster distance. The quantum efficiency for FRET is given by:

$$\begin{aligned} \zeta_{th} &= \frac{k}{1+k} \\ &= \frac{(\pi N_A R_o^6/2)}{r^4 + (\pi N_A R_o^6/2)} \end{aligned} \tag{S7}$$

By normalizing the numerator and denominator with respect to initial distance between fluorophores, $r_\mathrm{i}$, we obtain:

$$\begin{aligned}\zeta_\mathrm{th} &= \frac{\frac{\left(\pi N_\mathrm{A} R_\mathrm{o}^6/2\right)}{r_\mathrm{i}^4}}{\left(\frac{r}{r_\mathrm{i}}\right)^4 + \frac{(\pi N_\mathrm{A} R_\mathrm{o}^6/2)}{r_\mathrm{i}^4}} \\ &= \frac{c}{\left(\frac{r}{r_\mathrm{i}}\right)^4 + c}\end{aligned} \qquad [S8]$$

where $c = \pi N_\mathrm{A} R_\mathrm{o}^6/(2r_\mathrm{i}^4)$. Because of non-dimensionalization with respect to initial distance, $r_\mathrm{i}$, we do not use explicit values of $R_\mathrm{o}$ and $N_\mathrm{A}$ for calculations. We calculate $c$ by equating experimental FRET efficiency, $\zeta_\mathrm{exp}$ with theoretical fret efficiency, $\zeta_\mathrm{th}$ closest to saturation ($\psi = 0$) such that $r = r_\mathrm{i}$.

Upon substituting the Eq. (S3) in Eq. (S8), we obtain the following relation between $\zeta_\mathrm{th}$ and $\phi$:

$$\zeta_\mathrm{th} = \frac{c}{\left(\frac{\phi_\mathrm{o}}{\phi}\right)^{4/3} + c}. \qquad [S9]$$

Combining the Flory-Rehner model (Eq. (S1)) relation between water potential ($\psi$) and volume fraction of polymer ($\phi$) with Eq. (S9), we obtain a one parameter model that relates $\zeta_\mathrm{th}$ with $\psi$:

$$\zeta_\mathrm{th} = f(\psi, c). \qquad [S10]$$

The single point calibration needed to generate a curve to predict $\zeta_\mathrm{th}$ for the range of interest of water potential, simplifies the calibration of $\zeta_\mathrm{th}$ vs. $\psi$ for each new batch of AquaDust nanoparticles.

***B.3. Wolber FRET Model (Model for uniform distribution of donor and acceptor fluorophores in infinite plane).*** As seen from the distribution of AquaDust on the leaf cell surfaces (Fig. 2 - main text), fluorophores can be assumed to be randomly distributed and spaced at ∼nm scale on a plane of a large $\mu$m scale cell surface. Accordingly, we consider the analytical solution presented by Wolber et al.(12) to model the FRET for donor and acceptor fluorophores randomly distributed on an infinite plane:

$$\begin{aligned}\zeta_{\text{th}}(\text{Wolber}) &= \sum_{j=0}^{\infty}\left[\frac{\left(-\pi\Gamma\left(2/3\right)R_{\text{o}}^2 n\right)^j \Gamma\left(j/3+1\right)}{j!}\right] \\ &= \sum_{j=0}^{\infty}\left[\frac{\left(-\pi\Gamma\left(2/3\right)R_{\text{o}}^2 n_{\text{o}}\left(\frac{n}{n_{\text{o}}}\right)\right)^j \Gamma\left(j/3+1\right)}{j!}\right] \\ &= \sum_{j=0}^{\infty}\left[\frac{\left(-\pi\Gamma\left(2/3\right)R_{\text{o}}^2 n_{\text{o}}\left(\frac{\phi}{\phi_{\text{o}}}\right)^{2/3}\right)^j \Gamma\left(j/3+1\right)}{j!}\right] \\ &= \sum_{j=0}^{\infty}\left[\frac{\left(-\pi\Gamma\left(2/3\right)c''\left(\frac{\phi}{\phi_{\text{o}}}\right)^{2/3}\right)^j \Gamma\left(j/3+1\right)}{j!}\right]\end{aligned} \quad [\text{S11}]$$

where $c'' = R_{\text{o}}^2 n_{\text{o}}$ is calculated by equating $\zeta_{\text{th}}(\text{Wolber})$ with the experimental FRET Efficiency, $\zeta_{\text{exp}}$ closest to saturation ($\psi_{\text{exp}} = 0$), and $\Gamma(x)$ is a gamma function evaluated at $x$. We consider first 400 terms as an approximation to calculate $\zeta_{\text{th}}$ from this infinite series.

**C. Design optimization for AquaDust response.** Our design objective is to maximize

$$\zeta_{\text{diff}} = \zeta_{\text{th}}(\psi = -3\,\text{MPa}) - \zeta_{\text{th}}(\psi = -0.1\,\text{MPa}) \quad [\text{S12}]$$

with the following constraints: (i) Percentage monomer concentration $\%T$ (w/v) $(= w_{\text{mon}} \times 100)$ should be greater than 2% (= 0.02 gm monomer/1 $\text{cm}^3$ solution ×100) for gelation (13), (ii) Relative cross-linker concentration $\%CL(= w_{\text{cl}}/w_{\text{mon}} \times 100)$ should be greater than 0.5% for gelation for the minimum value of $\%T$ (13), (iii) $\%CL$ should be less than 4% for synthesis of ideal-network gel and to avoid formation of heterogeneities or a clustered gel network (14).

We solve for $\zeta_{\text{diff}}$ using Eq. (S10) with the following values for different parameters: (i) We assume a constant Flory-Chi parameter, $\chi = 0.48$ (15, 16) as reported in literature for acrylamide gels (values range from $0.46 - 0.49$) with similar composition. We also found that we can fit the evolution of the volume of macroscopic polyacrylamide gel with constant value of $\chi = 0.48$ (described later in Sec. S3, Fig. S3C). (ii) We vary $\%CL$ from 1% to 3% and $\%T$ from 3% to 15% as prescribed by the constraints. (iii) We use three different values of fit parameter, $c = 0.1, 0.2$ and $0.3$ as shown in Fig. S1. As discussed before, $c$ is a function of the concentration of conjugated dye.

Fig. S1 shows that $\zeta_{\text{diff}}$ has a weak/negligible dependence on $\%CL$ in the bounded range of $1\% - 3\%$ ($\zeta_{\text{diff}}$ ($\%CL = 1\%$) - $\zeta_{\text{diff}}$ ($\%CL = 3\%$) <0.02). However, $\zeta_{\text{diff}}$ is a strong non-monotonic function of $\%T$ and attains maxima at different values of $\%T$ for different values of $c$. We chose a fixed dye concentration for AquaDust synthesis (correspondingly, fit parameter $c = 0.2$, see Sec. S4 J, Fig. S10), we chose $\%T$ to be 6.1% and $\%CL$ to be 3% for maximum $\zeta_{\text{diff}}$. Table S1 provides the information on the composition of polymer matrix and parameters used in bulk gel and nanoparticulate AquaDust synthesized as in Sec. S3 A and Sec. S4 A respectively.

**D. Comparing alternative models for FRET interaction.** We compared the theoretical prediction from the dipole-dipole, dipole-plane and Wolber FRET model (as described in Sec. S2 B) against the experimental values of water potential measured using AquaDust (Fig. 3 - main text) in Figure S2. These models share one and the same adjustable parameter (variable labeled as ($c$ in Eq. (S9), $c'$ in Eq. (S5) and $c''$ in Eq. (S11)) which is a function of initial dye separation. The fit parameter ($c$ in Eq. (S9), $c'$ in Eq. (S5) and $c''$ in Eq. (S11)) in each of these models is obtained by forcing $\zeta_{\mathrm{th}} = \zeta_{\mathrm{exp}}$ to the experimental data closest to saturation (here, $\psi_{\mathrm{PC}}^{\mathrm{leaf}} = -0.08$ MPa). In physical terms, the resulting value of $c = c' = 0.217$ corresponds to $r_{\mathrm{i}} = 1.3R_{\mathrm{o}}$. It is clear from Fig. S2 that the dipole-plane FRET model captures the AquaDust response in leaves, while the Wolber FRET model under-predicts the FRET Efficiency and dipole-dipole FRET model over-predicts the FRET Efficiency.

Since AquaDust contains randomly distributed fluorophores, it is unlikely that the donor interacts with a single acceptor fluorophore, as assumed by the dipole-dipole model. In dipole-dipole model, the FRET efficiency is more sensitive than the dipole-plane model to the distance between the fluorophores; this sensitivity possibly causes the over-prediction of FRET Efficiency with decreasing water potential observed in Fig. S2 (pink-dashed curve). On the other hand, the Wolber FRET model is associated with multi-fluorophore interaction among dyes confined in a plane; potential 3D interactions are not taken into account in this model, resulting in possible under-prediction of FRET Efficiency observed in Fig. S2 (blue-dashed curve). The dipole-plane FRET model incorporates the single donor-multi acceptor interaction in different planes. This scenario appears to be most descriptive of the distribution of fluorophores and their interaction in the AquaDust.

### S3. Synthesis and characterization of the macroscopic gels

We synthesized macroscopic bulk gels to characterize gel swelling and FRET as a function of the water potential of its environment, $\psi^{\mathrm{env}}$, guided by theoretical consideration (Sec. S2, Fig. S1). We consider composition of monomer and cross-linker to maximize the measurement sensitivity and resolution between saturation (water potential, $\psi^{\mathrm{env}} = 0$, water activity, $a_{\mathrm{w}} = 1$) to mild undersaturation ($\psi^{\mathrm{env}} = -3$ MPa, $a_{\mathrm{w}} \approx 0.98$ at temperature, $T = 25$°C ); this range is typical of water potential in leaves, and the choice of the monomers is guided towards minimizing the response of the gels to pH and temperature (Fig. S3, Table S1).

**A. Bulk gel synthesis.** We synthesized bulk gels corresponding to the composition adopted for the studies presented in this paper (designed for $0 < \psi < -3$ MPa, Table S1, Fig. S3). Fig. S4 shows the chemical constituents of the gel matrix used for bulk gel synthesis here (same for AquaDust synthesis described later (Sec. S4)). Bulk cylindrical polyacrylamide gel were synthesized as follows: 5.5%(w/v) AAm (275 mg), 0.18%(w/v) BisAAm (9 mg), 0.6%(w/v) APMA (30 mg) in 5 ml were stirred and sonicated in 100 mM phosphate-buffered saline (PBS) (pH 7.4) buffered solution (all PBS is in water); this solution was degassed in vacuum for 20 minutes to remove dissolved oxygen; 0.5 mg donor dye AF488 and 1 mg acceptor dye AF568 were added to the solution and stirred for 1 minute (Note: different FRET pair is used for synthesizing AquaDust, later in Sec. S4, compared to the macroscopic gel here due to cost considerations); polymerization reaction was triggered by adding initiator, 10 mg APS, and accelerator, 50 $\mu l$ TEMED; the solution was transferred prior to gelation (in <3 minutes after adding the initiator and accelerator) into 3 mm inner diameter glass tubes using a syringe (Hamilton Gastight Syringe, 5ml). The tube served as a mold to define a cylindrical gel. The solutions were allowed to polymerize for 24 hours at room temperature. Intact gel was removed from the tube by blowing air from one side of the tube. It was then placed in MilliQ water and stirred for 1 day to remove the unreacted components and allowed it to come to saturated equilibrium.

**B. Instrumentation for ex-situ bulk gel fluorescence measurement.** The swollen gel was transferred in a temperature controlled vacuum chamber (Fig. S3A) on a dichlorodimethylsilane coated silicon wafer (to minimize adhesion of gel with the substrate) with a control on water vapor activity, $a_{\mathrm{w}}$ within ±0.1% (we have described this vacuum stage previously (17)). The gel was allowed to come to equilibrium at a set vapor activity. The color images (Fig. S3B) were captured using CanonShot Power G6 7MP camera attached to the Leica MZ FLIII stereo-microscope using a GFP Plus longpass filter (Excitation: $480 \pm 20$ nm, Emission: $> 510$ nm). A mercury lamp light source (Leica EL6000) was used as source for illumination, with a narrow-band optical filter (470 nm-500 nm) to select the excitation light wavelength. The reflected light captured by the central fiber (QR600-7-UV-125F, Premium 600 micron Reflection Probe, Ocean Optics Inc.) was passed through a long-pass cutoff filter ($> 510$ nm) and sent to the spectrometer (Ocean Optics Inc., ST2000); the collection filter helped avoid the saturation of detector with excitation light. The spectra (Fig. S3B) were saved using OceanView software operating with an integration time of 0.1 sec averaged 3 times.

***B.1. Calculating experimental FRET Efficiency.*** We determined the relative contributions of the donor and acceptor emissions ($x$ and $y$) by spectrally decomposing the emission spectra, $Em_{\mathrm{exp}}$:

$$\begin{bmatrix} Em_{\mathrm{Donor}} & Em_{\mathrm{Acceptor}} \end{bmatrix} \begin{bmatrix} x \\ y \end{bmatrix} = Em_{\mathrm{exp}} \qquad \text{[S13]}$$

where $Em_{\mathrm{Donor}}$ and $Em_{\mathrm{Acceptor}}$ are the normalized emission spectra of donor and acceptor as supplied by the chemical vendor. Experimental relative FRET Efficiency was calculated as the following ratio:

$$\zeta_{\text{exp}} = \frac{y}{x+y} \quad \text{[S14]}$$

**C. Bulk gel behavior with decreasing water potential and comparison with theoretical model.** At fixed temperature and gel composition, Eq. (S1) provides a one-to-one monotonic inverse relationship between matrix concentration, $\phi$, and $\psi^{\text{matric,gel}}$. In turn, if we allow the gel to come to equilibrium with an environment at $\psi^{\text{env}}$ such that $\psi^{\text{matric,gel}} = \psi^{\text{env}}$, we expect $\phi$ to vary with $\psi^{\text{env}}$ (Eq. (S1)). As $\psi^{\text{env}}$ decreases, water leaves the matrix such that it shrinks (decreasing volume) and the concentration of polymer, $\phi$, increases. In Fig. S3 B-C, we see a $\sim$ 13 fold decrease in volume ($V_o/V$ decreases) and increase in $\phi$ as $\psi^{\text{env}}$ goes from 0 MPa to -4 MPa. We note good agreement with Eq. (S1) with $n_o$ and $\phi_o$ set by the parameters of the synthesis (Sec. S2, Table S1) and a constant value of Flory temperature, $\theta = 277$K at $T = 288.2$K (equivalently, interaction parameter, $\chi = \theta/2T = 0.48$); these values of $\theta$ and $\chi$ are similar to the interaction parameters reported for gels with similar composition (16). We here consider fresh gels (curing time $<$ 24 hours and experimentation in less than 2 days). Anomalies in the interaction parameter have been reported for bulk gels allowed to cure for more than 6 days (18). We did not observe such anomalies with our work with AquaDust (nanogels) as we used each batch of AquaDust for $\sim$ 30 days.

We saw a qualitative change in the fluorescence spectrum of the gel with the swelling in the color change in Fig. S3B and in the rise in the relative intensity of Alexa Fluor 568 at $\sim$ 605 nm in Fig. S3B. We captured the fluorescence emission spectra of these gels (Fig. S3B) and extracted the value of experimental FRET Efficiency ($\zeta_{\text{exp}}$) using spectral decomposition (Sec. S3B.1) of the fluorescence emission spectra (Fig. S3D). Upon choosing dipole-plane model for FRET interaction, Eq. (S1) and Eq. (S8) combined has one free parameter, $c$ to fit the experimental data. We find that the theoretical FRET Efficiency, $\zeta_{\text{th}}$ using a constant value of $c$ ($c = 0.028$ in Eq. (S8)) combined with the model for the volume change of the gel (Eq. (S1)) is consistent with the evolution of $\zeta_{\text{exp}}$ with $\psi^{\text{env}}$ as shown in Fig. S3D. See Sec. S2B,D for discussion of the choice of FRET model.

We extend this understanding to develop nanogels with the same gel composition (AquaDust) as a tool for point-wise measurement of the water potential in leaves.

### S4. Synthesis and characterization of AquaDust

**A. Nanoparticle synthesis.** Nanoparticles were synthesized using inverse micro-emulsion procedure as described in literature (19, 20). Briefly, the aqueous polymerization solution contained 5.5%(w/v) AAm (110 mg), 0.18%(w/v) BisAAm (3.6 mg), 0.6%(w/v) APMA (12 mg) in 2 ml, 100 mM phosphate-buffered saline (PBS) (pH 7.4) buffer and was sonicated prior to use. Dyes were incorporated after particle synthesis (see next subsection). Hexane was deoxygenated by purging it with nitrogen for 15 minutes. 42 ml of deoxygenated hexane, 1.4 gm AOT and 2.88 ml Brij30 were stirred using a magnetic stirrer in a 100 ml round-bottom flask under a nitrogen atmosphere at room temperature, to which 2 ml of polymerization solution was added. The emulsion was sonicated for 4 minutes and stirred at 400 rpm for 4 minutes to form the microemulsion. To the microemulsion, 10 mg (dissolved in 50 $\mu l$ water) APS and 25 $\mu$l TEMED was added to initiate free-radical polymerization and the solution was stirred at 400 rpm for 2 hours. After 2 hours, hexane was removed using rotary evaporation at a pressure of 300 mbar at 40°C . The resulting suspension of particles was washed five times by suspending particles in 40 ml 100% ethanol (acts as anti-solvent) and precipitating using centifugation (8000 RPM $\equiv$ 7500 RCF, 20°C , 5 minutes).

**B. Dye functionalization.** Dried nanoparticles ($\sim$ 50 mg) were re-suspended in 5 ml 0.1M Sodium bicarbonate/Sodium Carbonate buffer (pH 8.5) by ultra-sonicating it for 10 minutes. To the suspension, 5 mg of NHS-Ester functionalized Oregon Green 488 was dissolved in 500 $\mu l$ of anhydrous DMF and 10 mg of NHS-Rhodamine was dissolved in 1000 $\mu l$ of anhydrous DMF. The solution was stirred at room temperature using a magnetic stirrer at 500 rpm for 3 hours. The conjugated nanoparticles were purified using MWCO 10000 Slide-A-Lyzer Dialysis kit against Milli-Q water for 3 days; the buffer was exchanged six times during the process. Once purified, the nanoparticles were characterized for size and concentration. The chemical composition of AquaDust is shown in Figure S4.

**C. AquaDust Size and Concentration.** To assess the size distribution of AquaDust, 0.7 mg of dry AquaDust nanoparticles were re-suspended in 2 ml Milli-Q water by sonicating it for 10 minutes. Particle size distribution was measured using a Malvern Nano ZS Zetasizer in 173° back-scatter configuration. Figure S5 shows the size distribution of a typical sample of AquaDust: Size distribution with respect to scattering intensity had a mean diameter of 80 nm with standard deviation of 30 nm; the distribution with respect to volume had a mean diameter of 56 nm with a standard deviation of 23 nm; and the distribution with respect to number had a mean diameter of 42 nm with a standard deviation of 13 nm, with a polydispersity index of 0.18.

To assess the concentration of particles in the suspension infiltrated into leaves, AquaDust suspension was diluted 10 fold in MilliQ water (i.e., 0.7 mg AquaDust nanoparticles in 20 ml Milli-Q water) and the concentration was evaluated using Malvern NS300 NanoSight. Concentration of particles in AquaDust suspension was found to be $6.6 \times 10^8 \pm 1.9 \times 10^8$ (Mean $\pm$ Std. Error) particles/ml. This concentration, with integration time of 1.2 sec (see next section for details on instrumentation), allowed AquaDust fluorescent intensity to be at least 10 fold larger than leaf autofluorescence.

**D. Microscopy for AquaDust distribution.**

***D.1. Top-view imaging (Fig. 2 B,C - main text).*** Maize leaf was infiltrated with AquaDust a day prior to imaging. A cut-out of the maize leaf ($\sim 2$ cm $\times \sim 5$ cm ($x \times y$), Fig. 2A - main text) infiltrated with AquaDust was mounted on a glass slide using a double-sided tape, placed on Zeiss LSM880 Confocal Upright Microscope, and imaged using W Plan-Apochromat 20x/1.0 water immersion objective.

***D.2. Cross-section imaging, (Fig. S6).*** A cut-out of a maize leaf infiltrated with AquaDust was placed in 20% (w/v) Agarose (Neta Scientific Inc., Garde-Biotech) (heated to 60°C ) in a rectangular mold and allowed to solidify. The embedded tissue was sectioned along the xz-pane using KD-400 vibrating blade microtome into 200 $\mu$m-thick slices (Fig. S6). The tissue sections were imaged using Zeiss LSM710 confocal microscope with Plan-Apochromat lens 63x/1.40 Water mode with tissue placed between two cover slips.

**E. Photosynthesis and gas exchange parameters in region with and without AquaDust.** To test if the process of using AquaDust to measure leaf water potential would affect leaf physiological functions, we compared gas exchange rates between maize leaf area with and without infiltration of AquaDust. The gas exchange processes tracked were transpiration rate ($E$), water vapor conductance ($G_{H_2O}$) and assimilation rate ($A$). Five B97 maize plants at V9 to V10 leaf growth stages were used in the experiment. These plants were grown in a Cornell University Guterman lab growth chamber at 14 h light and 10 h dark. We infiltrated the middle part of Leaf 7 with AquaDust one side of the mid-rib. After 24 hours of infiltration, the portable gas exchange fluorescence system GFS-3000, the standard measuring head 3010-S, and light source of LED-Array/PAM-Fluorometer 3055-FL (Walz, Effeltrich, Germany) were used to measure these physiological parameters on the infiltrated leaf area and non-infiltrated leaf area that was on the other side of the mid rib in the same leaf. The measurements were performed with a chamber $CO_2$ concentration of 400 ppm, photosynthetic quantum flux density of 500 $\mu$mol/(m$^2$.s), chamber relative humidity at 50%, air flow rate of 750 $\mu$mol/s, and cuvette temperature of 25°C . Paired sample two-tailed t-test was performed to test if there were significant differences of physiological traits between infiltrated (AquaDust, Table S2) and non-infiltrated (Control, Table S2) leaf area of the same leaf.

A second experiment was conducted to test if AquaDust, as a chemical per se, could influence leaf physiological parameters. Four B97 maize plants at V9 to V10 stages were used. AquaDust was infiltrated in middle part of leaf 6 on one side of the mid rib, while water was infiltrated on the other side. We also used GFS-3000 to measure the gas exchange rates of the maize leaf area infiltrated with AquaDust and water. Paired sample two-tailed t-test was performed to test if there are significant differences between AquaDust and water infiltrated areas in the same leaf for the measured physiological parameters.

For both experiments, the p-value of paired sample two-tailed t-tests for $E$, $G_{H_2O}$ and $A$ were larger than 0.05. We conclude that the infiltration of AquaDust caused no significant difference in these processes relative to non-infiltrated or water-infiltrated leaf areas.

**F. AquaDust response to temperature.** We calibrated response of AquaDust to temperature under saturation conditions ($\psi^{\mathrm{env}} = 0$) using the setup as described earlier (17) (Fig. S7A, Sec. S3B). We pipetted AquaDust on Nylon (Plain, 0.8 micron pore diameter, Fisher Scientific Inc.) and placed the sample on a plasma-cleaned silicon wafer in a vacuum chamber equipped with an optical window which allows collection of the fluorescence spectra using reflection probe; the sample rested on the bottom surface of the chamber which was thermostated at temperature, $T$, using a water bath and temperature of the stage was measured using a thermocouple. We decrease $T$ in steps (Fig. S7B) while keeping the vapor pressure in the chamber at saturation vapor pressure for the imposed $T$. We estimated the uncertainty on the temperature to be $< 0.1^{\mathrm{o}}$C from the measured fluctuations of $T$ during the experiments. Fig. S7C shows the fluorescence spectra from AquaDust with changing temperature as indicated in Fig.S7B. Based on water potential calibration at room temperature ($\sim 24^{\mathrm{o}}$C ), we calculated the value of water potential based on theoretical calibration model as plotted in Fig. S7D.

Response of polyacrylamide gel to temperature is usually small compared to other homopolymers (21, 22). Okano et al. (21) attributed the slight increase in volume of the gel with increasing temperature to the weakened hydrogen bonding interactions. This swelling is consistent with slight decrease in emission from second dye (and experimental water potential obtained from AquaDust $\psi_{\mathrm{AQD}}$, Figure S7 C,D) with increasing temperature. We observed an outlier from the trend. Such small variation in water potential can result from varying concentration of AquaDust nanoparticles or errors associated with background correction. Nonetheless, all inferred variations in water potential with changes in temperature ($\sim 0.006$ MPa; Fig. S7D) were very small compared to the uncertainty associated with AquaDust measurement of $\pm 0.14$ MPa (refer Sec. S4 J).

**G. AquaDust response to pH.** AquaDust was suspended in 0.1 M sodium acetate buffers (pH 5-7) and 0.1 M sodium carbonate/bicarbonate buffers (pH 8-11) and pH was monitored using pH strip (BDH pH test strip, VWR Inc., with a resolution of 0.3 difference in pH) with fixed concentration of AquaDust (as used for AquaDust infiltration in leaves, Sec. S4 C) and measured the emission spectra of AquaDust using the spectrofluorometer (PTI Quantamaster 8000) with an integration time of 50 ms averaged over 5 readings. Buffer solutions without AquaDust provided the emission spectra associated with background and was subtracted from the emission spectra of the AquaDust solutions. Fig. S8A shows the averaged fluorescence spectra from AquaDust as a function of pH. We observe that emission intensity from AquaDust decreases by $\sim 70\%$ of the maximum intensity with decreases in pH below 6 and is likely associated with decrease in emission from the donor dye Oregon Green 488 (6, 23).

Fig. S8B shows the fluorescence spectra normalized with respect to emission peak of donor and Fig. S8C shows corresponding values of FRET efficiency as calculated from these spectra. FRET efficiency remains constant for pH ranging from 6-11 with slight increase in pH below pH 6. As seen from Fig. S8C, the corresponding water potential as calculated from $\zeta_{exp}$ and the theoretical model (Eq. (S10)) is reasonably within the AquaDust resolution of ±0.14 MPa (refer Sec. S4 J), yet, could be one of the sources of error in measurement of FRET Efficiency. This source of error could be minimized by choosing dyes that are insensitive to pH such as Alexa Fluor 488 and Alexa Fluor 555 as donor dye and acceptor dye respectively.

**H. Instrumentation for *in-planta* measurement.** Fig. 3A (main text) shows the sketch of the setup used for in-situ measurement. Briefly, a mercury lamp light source (Leica EL6000) was used as source for illumination, with a narrow-band optical filter (470 nm-500 nm) to select the excitation light wavelength used to excite AquaDust at the site of interrogation in leaf. The excitation light is directed using a reflection probe (QR600-7-UV-125F, Premium 600 micron Reflection Probe, Ocean Optics Inc.) fitted in a collimator (74 VIS Fiber Optic Collimator, Ocean Optics Inc.) and positioned on the leaf using a leaf clamp (Fig. S9). This clamp serves to minimize scattered/stray light on measurement. Direct contact of fiber bundles does not occur, for example, in fully shaded (no stray light) environment, the fiber bundle could be used to collect fluorescence signal from a distance of several mm. The reflected light captured by the reflection probe was passed through long-pass cutoff filter ($>$ 510 nm) and sent to a spectrometer (Ocean optics Inc., ST2000). The spectra were saved using OceanView software operating with an integration time of 1.2 sec averaged 3 times. The integration time was chosen such that the intensity of fluorescence collected from AquaDust was less than 70% of the saturation limit of the spectrometer. The fluorescence spectra from the leaf not infiltrated with AquaDust was saved as background signal and subtracted from the spectra collected from AquaDust infiltrated leaf. Depending on the length of the infiltration of AquaDust, a minimum of 3 spectra and a maximum of 6 spectra were recorded from the same infiltration site by moving the optical probe across the infiltration zone. No distinction was made in the data collected from the upstream/downstream of the AquaDust-infiltration site. Spectra from locations within $<$ 1 cm of site of infiltration were not used (see Sec. S4 K). Experimental relative FRET efficiency, $\zeta_{exp}$, were calculated as described previously in Sec. S3 B.1. Mean FRET Efficiency was calculated by averaging all the recorded 3 to 6 spectra. Table S3 contains the numerical values of data in Fig. 3C (main text).

**I. *In planta* calibration, Fig. 3 (main text).** A total of 36 maize plants (inbred line B97) at the V4 to V5 stages were used in the experiment presented in Fig. 3 (main text) with 18 plants under a water-limited treatment (no water applied) and 18 plants under a well-watered (control) treatment. These 36 plants were grown in a greenhouse under long-day lighting conditions (16h light/8h dark cycle) provided by 400 watt high pressure sodium lamps. Each maize plant was grown in 15 cm pot (1L of volume) and filled with an artificial soil mix (Cornell mix: 2 bales (0.11 $m^3$) peat, 2 bags (0.22 $m^3$) vermiculite, 1 bag (0.11 $m^3$) perlite, 1.8 kg calcium sulfate, 0.11 kg Aquagrow 2000 (wetting agent), 1.8 kg 10-5-10 Media Mix, 2.3 kg limestone).

We define the tip of the leaf as one-third of the total leaf length from the tip end of the leaf. For leaf 4/5, the tip region is approximately 25 cm, while for leaf 6/7, the tip region is approximately 30 cm. AquaDust was infiltrated in the tip region of the attached leaves 4/5 and 6/7 in 3 well-watered and 3 water-limited plants at 10 am on Day 0. (Note: For our experiments, each batch of AquaDust was synthesized with the quantities as described in Sec. S4A, and it allowed for $\sim$ 200 $\psi^{\text{leaf}}$ measurements.) Depending on the length of the infiltration of AquaDust, a minimum of 3 spectra and a maximum of 6 spectra were recorded from each replicate on Day 1 at pre-dawn (5.00 A.M. - 6.00 A.M. EST) and mid-day (11.00 A.M. - 2.00 P.M. EST). Mean FRET Efficiency is calculated by averaging all the recorded 3 to 6 spectra and the range for FRET Efficiency correspond to the minimum and maximum values of FRET Efficiency of all the spectra. Each AquaDust measurement takes $<$ 30 seconds. Table S3 contains the numerical values of minimum, mean, and maximum value of FRET Efficiency from each replicate (plant) and is plotted in Fig. 3C (main text).

Right after the AquaDust spectra were recorded for all three replicates (plants), pressure chamber measurements of water potential ($\psi_{\text{PC}}^{\text{leaf}}$) were taken from the tip region of leaf 4/5 and leaf 6/7. The uninfiltrated side across the midrib of tip region of the maize leaf was severed (not including the midrib) and then immediately wrapped with a plastic bag. The wrapped leaf section was quickly inserted into the pressure chamber (Model 615, PMS Instrument Company, Albany, Oregon, USA) with the cut end protruding through the chamber seal. Chamber pressure was slowly increased until the endpoint defined by water just appearing at the cut surface. $\psi_{\text{PC}}^{\text{leaf}}$ was considered to be equal to the negative value of the pressure applied when water appeared. The pressure bomb measurement from a particular plant from which AquaDust measurements were taken served as the mean $\psi_{\text{PC}}^{\text{leaf}}$ for that particular plant; the range was calculated from the pressure chamber measurements of three replicates (plotted in Fig. 3C - main text). This process was repeated each day for the next set of 3 well-watered and 3 water-limited plants for a 7 day period with no water applied to the water-limited plants. Table S3 contains the numerical values of minimum, mean, and maximum value of pressure chamber water potential measurements using from all three replicates (plotted in Fig. 3C - main text).

**J. Calculation of uncertainties from AquaDust results.** Here, we elaborate on the methods used to calculate uncertainties on the calibration fit (Fig. S10A) and the confidence interval on $\psi_{\text{AQD}}^{\text{leaf}}$.

*Goodness of model with fitting parameter $c$*: As described in Sec. S2B, we used a single-point calibration to predict $\zeta_{\text{th}}(\psi)$ using Eq. (S10) (Flory-Rehner theory for gel swelling and dipole-plane

FRET model for fluorophore interactions), where we found the value of $c$ by equating the water potential ($\psi^{\text{leaf}}_{\text{AQD}}$) calculated from the mean relative fret efficiency ($\zeta_{\text{exp}}$, Fig. 3C - main text) with that of the water potential from the pressure chamber ($\psi^{\text{leaf}}_{\text{PC}}$) closest to saturation (here, at $\psi^{\text{leaf}}_{\text{PC}} = -0.08$ MPa). We obtained $c = 0.217$ and plotted the corresponding $\zeta_{\text{th}}$ as shown in Fig. S10A. We plotted the corresponding residual (difference between $\psi_{\text{AQD}}$ and $\psi_{\text{PC}}$) in Figure S10B. First, we note that there is no obvious trend in the residuals; this lack of a trend in the deviation of the data from the model supports the appropriateness of the model. From Figure S10B, we found that averaged over all the readings, the absolute mean difference between the water potential obtained from AquaDust and pressure chamber was 0.018 MPa with a standard deviation of 0.067 MPa.

*Confidence interval on the fit parameter c:* We explain the process used to find the uncertainty associated with the parameter $c$. We obtained the experimental range of water potential ($\psi_{\text{AQD}}$) from the range of AquaDust FRET Efficiency using the theoretical prediction curve as shown in Figure S10A (red-dashed lines). The value of this range divided by 2 is plotted as uncertainty in Figure S10C. We found the value of $c$ (Eq. (S10)) such that the absolute sum of the difference between the minimum experimental $\psi_{\text{AQD}}$ (left end of the red dashed line in Figure S10A) and the water potential from the pressure chamber ($\psi_{\text{PC}}$) is minimized. Consequently, we obtained $c = 0.199$ and plotted the theoretical value of FRET Efficiency $\zeta_{\text{th}}$ from Eq. (S10) as shown in Figure S10A. Similarly, we found the value of $c$ (Eq. (S10)) such that the absolute sum of the difference between the maximum experimental $\psi_{\text{AQD}}$ (right end of the red dashed line in Figure S10A) and the water potential from the pressure chamber ($\psi_{\text{PC}}$) is minimized. Consequently, we obtained $c = 0.231$ and plotted the theoretical prediction from Eq. (S10) as shown in Figure S10A. We found that the value of fitting parameter, $c$, can range from 0.199-0.231.

*Inferences from model:* From Figure S10C, the average of range of uncertainty was ±0.08 MPa and a standard deviation of ±0.03 MPa; we can resolve the water potential from AquaDust with 95% confidence interval (= mean of the range added to 1.96 times the standard deviation) of ±0.14 MPa.

**K. Characterization of mechanical damage during infiltration.** To evaluate the effect of mechanical damage to the cuticle caused during infiltration on the local water potential, we measured AquaDust spectra as function of distance from spot of infiltration (Figure S11). We calculated the relative FRET efficiency and corresponding water potential, and observed that the effect on FRET efficiency was limited to ±3 mm from the injection site. The FRET efficiency was uniform beyond this spatial effect. Accordingly, we ensured that the AquaDust measurement was taken at least 1 cm away from the site of infiltration.

**L. Response time to step change in water potential using a pressure chamber.** We evaluate the response time of AquaDust for step change in water potential by forcing a transpiring maize leaf to saturation using a pressure chamber (Fig. S12). Briefly, we sever a maize leaf infiltrated with AquaDust from one side of the mid- vein. We place the exposed side in a water-filled Bitran bag (Fisher Scientific Inc.) with the rest of the leaf sticking out of the rubber gasket. The gasket is allowed to seal the chamber and pressurized to 0.2 MPa. The leaf gets saturated instantaneously as water exudes from the leaf sticking out of the chamber. Spectra measurements were taken and corresponding theoretical water-potential ($\psi^{\mathrm{leaf}}_{\mathrm{AQD}}$) was calculated from the calibration curve (Eq. (S10)). We observed that $\psi^{\mathrm{leaf}}_{\mathrm{AQD}}$ returned to saturation in $\sim 1$ minute. This suggests AquaDust responds to changes in water potential in order of minutes.

**M. Calibration in the field.** We performed in-planta comparison of water potential measurement from AquaDust and Scholander pressure chamber in field-planted maize (Musgrave Research Farm, Location: 42°43′N, 76°39′W). For the Day 1 measurement (20 July 2018, 3.00 P.M. - 5.00 P.M. EST), AquaDust was infiltrated 1 day before (19 July 2018, 10.00 A.M - 12.00 P.M. EST) on maize plants at leaf growth stages V10-V12. For Day 2 measurements (22 July 2018, 11.00 A.M. - 1.00 P.M. EST), AquaDust was infiltrated 2 days before on 20 July 2018. We report these data of experimental relative FRET efficiency in Fig. S13 as a function of water potential measured using a pressure chamber for the same maize plant. We plot the theoretical FRET Efficiency, $\zeta_{\mathrm{th}}$ using a constant value of $c$ (here, for this batch of AquaDust, $c = 0.19$ in Eq. (S8) chosen such that $\zeta_{\mathrm{th}} = \zeta_{\mathrm{exp}}$ at $\psi^{\mathrm{leaf}}_{\mathrm{PC}} = -0.38$ MPa) combined with the model for the volume change of the gel (Eq. (S1)). Out of total 11 measurements on leaf tip, 3 measurements show large difference from theoretical FRET. Possible sources of error include:

1. Outliers (i) and (ii) (See Figure S13): The pressure bomb measurements from Day 1 were taken on Leaf 3 while the AquaDust measurements were taken from Leaf 4. For different plants, the pressure bomb readings ranged from -0.3 to -0.8 MPa for Leaf 3 on Day 1 and -0.5 to -0.8 MPa on Leaf 4 on Day 2. This large variability in pressure chamber water potential measurements suggests that the local water potential measured by AquaDust could be different from the water potential measured using a pressure chamber leading to these outliers. Difference in light conditions between Leaf 3 and Leaf 4 could be a potential source of the observed difference between AquaDust and pressure chamber.

2. Outlier (iii): On Day 2, measurements from AquaDust were taken from the same leaf as the measurements from pressure chamber. Qualitatively, large fluctuations in local water potential due to uneven cloud cover and aberrant wind conditions on Day 2 could have resulted in the observed difference. This measurement was taken after brief rain while weather transitioned from cloudy to sunny in a short period of time and could have caused rapid changes in leaf water potential.

### S5. Theoretical model for leaf water potential in a transpiring leaf

**A. Measuring gradients in water potential along a leaf, Fig. 4 (main text):.** Eight 'B97' maize plants at V9 to V10 leaf growth stages were used in the experiment, which were grown in Cornell University Guterman Lab growth chamber under a light condition of 12 h light (06.00 A.M. - 06.00 P.M. EST) and 12 h dark, with 4 plants under water limited treatment and 4 in well-watered treatment (control) with following growth conditions: the daytime temperature was set at 31°C and nighttime temperature at 22°C ; PAR measured at the canopy height averaged at 270 $\mu$mol photons $m^{-2}$ $s^{-1}$, and air relative humidity varied from $45 - 50\%$.

Four water-limited replicates of maize plants were infiltrated with AquaDust on leaf 7 on Day 0 of water-limited condition and were used for continuous measurement. Each leaf was divided into three regions, tip being the first one-third of the leaf from the leaf end, mid being the second one-third portion of the leaf and node being the last one-third region of the leaf connected to the stem. Two or three infiltrations of AquaDust were made in each region. Four well-watered replicates were infiltrated in the same manner and the response of AquaDust was recorded for 3 days. The mean and error in FRET Efficiency was calculated from all the spectra recorded for all the infiltrations in the given region. The expected water potential range is calculated from the experimental FRET Efficiency using the theoretical curve shown in Fig. 3C (main text). Transpiration rate ($E$) was measured using portable gas exchange fluorescence system GFS-3000, the standard measuring head 3010-S, and light source of LED-Array/PAM-Fluorometer 3055-FL (Walz, Effeltrich, Germany).

**B. Model considering only xylem resistance.** We calculate theoretical xylem water potential ($\psi_{\text{th}}^{\text{xyl}}$) using measured values of transpiration rate, $E = 4.2 \times 10^{-5} \pm 0.85 \times 10^{-5}$ kg/(m$^2$.s) (Mean ± Range), for both well-watered and water-limited case, i.e., uniform rate of loss of water per length of leaf (from the node towards the tip of the leaf, See SI - Sec. S1 D for details of model and measurements). We acknowledge that this assumption of constant rate of transpiration as a function of position and potential is an over simplification. We believe it is justified in this analysis, because: i) the measured variation in E is modest (±20%); ii) in the range of stresses ($\psi^{\text{leaf}} \geq -1.5$ MPa) we do not expect significant stomatal closure (24); and iii) as we show in Fig. 4D (main text), this simplified model can provide predictions that are quantitatively consistent with our measurements of stress.

We first consider a model with only xylem resistance for comparison with the model presented in Fig. 4C (main text). As depicted in the hydraulic circuit in Fig. S14A, we hypothesize that the xylem presents the limiting, $\psi$-dependent resistance to flow to the sites where we measure $\psi$ with AquaDust, and we neglect extra-vascular resistance. The flow, $J(z)$ [kg/s] through the leaf xylem decreases linearly due to constant transpiration, $E$ [kg/(m$^2$.s)], and can be approximated as follows (neglecting axial variation of width):

$$J(z) = wE(L - z), \tag{S15}$$

such that $J(z = L) = 0$, where $w$ [m] is the average width of the leaf and $L$ [m] is the length of the leaf. The water potential gradient in the leaf along the z-direction is calculated using Ohm's law formulation as shown in Fig. S14A. At a given location, $z$, on the leaf, the gradient in xylem water potential, $\psi_{\text{th}}^{\text{xyl}}$, resulting from the xylem resistance, $R_{\text{xyl}}(\psi)$, to the flow through the xylem, $J(z)$ is given as follows:

$$\frac{\partial \psi_{\text{th}}^{\text{xyl}}}{\partial z} = -\frac{R_{\text{xyl}}(\psi)}{wL^2} J(z). \tag{S16}$$

For $R_{\text{xyl}}(\psi)$, we fit a three-parameter logistic function to the vulnerability curve obtained from Li et al. based on a centrifugal method in maize (25) (similar values for xylem resistance are obtained using other methods such as high-pressure flow meter (26)) such that,

$$R_{\text{xyl}}(\psi) = R_{\text{xyl}}^{\text{min}} \left(1 + \left(\frac{\psi_{\text{th}}^{\text{xyl}}}{\psi_{\text{th,50\%}}^{\text{xyl}}}\right)^a\right), \tag{S17}$$

where $R_{\text{xyl}}^{\text{min}} (= 3.47 \times 10^3 \text{ m}^2\text{.s.MPa/kg})$ is the minimum axial resistance, $\psi_{\text{th,50\%}}^{\text{xyl}} = -1.58$ MPa is the value of xylem pressure at which resistance increases by a factor of 2 (or conductance drops to 50% of its saturated value), and the logistic growth rate, $a = 4.5$ (25). We plotted xylem conductance, $K_{\text{xyl}} = 1/R_{\text{xyl}}$ in Fig. S14B.I as a function of $\psi$. We used average value of transpiration rate, $E$ $(= 4.2 \times 10^{-5} \text{ kg/(m}^2\text{.s))}$, measured on six different plants at ambient PAR ($E$ ranged from $3.4 \times 10^{-5}$ kg/(m$^2$.s) to $5.1 \times 10^{-5}$ kg/(m$^2$.s), as measured using GFS3000, Walz Inc. and averaged over 6 measurements on 3 well-watered and 3 water-limited plants on Day 3, see details of growth conditions and measurement in Sec. S5 A). Using Eq. (S15) for flux, $J(z)$, we solve the combined Eq. (S15), (S16) and (S17) numerically with the boundary condition such that the value of water potential $\psi_{\text{th}}^{\text{xyl}} = \psi_{\text{AQD}}^{\text{leaf}}$ at the node ($z = L/6$) and plot it as shown in Fig. S14B.II-IV.

We observed large difference in the theoretical prediction of $\psi_{\text{th}}^{\text{xyl}}$ and the experimental value of $\psi_{\text{AQD}}^{\text{leaf}}$ (Fig. S14B.II-IV) at both mid and tip of the maize leaf. This lack of agreement suggests that either the values used for $R_{\text{xyl}}$ are incorrect or additional resistances separate the site of $\psi_{\text{AQD}}^{\text{leaf}}$ measurement from the xylem. We test the former by adjusting the parameters for $R_{\text{xyl}}$ (Eq. (S17)) to minimize the difference between $\psi_{\text{th}}^{\text{xyl}}$ with $\psi_{\text{AQD}}^{\text{leaf}}$ for WW plants. Specifically, we vary $R_{\text{xyl}}^{\text{min}}$ (ranging from $3.47 \times 10^3$ m$^2$.s.MPa/kg to $3.47 \times 10^4$ m$^2$.s.MPa/kg) and $\psi_{\text{th,50\%}}^{\text{xyl}}$ (ranging from $-1.58$ MPa to $-0.5$ MPa) such that,

$$\sum_{\text{Day 1,2,3}} (\psi_{\text{AQD}}^{\text{leaf,mid}} - \psi_{\text{th}}^{\text{xyl}}(z = L/2)) + (\psi_{\text{AQD}}^{\text{leaf,tip}} - \psi_{\text{th}}^{\text{xyl}}(z = 5L/6)) \tag{S18}$$

is minimized, for different values of $a$ as shown in Fig. S14(C.I, D.I, E.I) and compare corresponding prediction for $\psi_{\mathrm{th}}^{\mathrm{xyl}}$ with $\psi_{\mathrm{AQD}}^{\mathrm{leaf}}$ in WW and WL plants as shown in Fig. S14(C.II-IV, D.II-IV, E.II-IV). We found that no set of parameters for $R_{\mathrm{xyl}}$ could predict $\psi_{\mathrm{AQD}}^{\mathrm{leaf}}(\pm$ range) in WL plants on all 3 days. For example, Fig. S14C.I, we found that minimum xylem resistance ($R_{\mathrm{xyl}}(\psi = 0)$) had to be $\sim$ 6-fold larger ($2 \times 10^4$ m$^2$.s.MPa/kg rather than $3.4 \times 10^3$ m$^2$.s.MPa/kg (25)) than the resistance reported in literature, and the value of $\psi_{\mathrm{th},50\%}^{\mathrm{xyl}}$ (the water potential inducing 2-fold increase in hydraulic resistance) had to be reduced $\sim$ 3 fold ($-0.6$ MPa rather than $-1.6$ MPa (25)), so that $\psi_{\mathrm{th}}^{\mathrm{xyl}}$ matched $\psi_{\mathrm{AQD}}^{\mathrm{leaf}}(\pm$ range) in Day 1 WL plants (Fig. S14C.I); for these values the corresponding prediction of $\psi_{\mathrm{th}}^{\mathrm{xyl}}$ did not match $\psi_{\mathrm{AQD}}^{\mathrm{leaf}}(\pm$ range) in WL plants on Days 2 and 3. This exercise excludes the simple model for leaf resistance with xylem as limiting resistance, and motivates addition of extra-vascular resistance ($R_{\mathrm{ox}}$) to xylem resistance and comparing prediction for outside-xylem water-potential ($\psi_{\mathrm{th}}^{\mathrm{ox}}$) with $\psi_{\mathrm{AQD}}^{\mathrm{leaf}}$ (Fig. 4C,D - main text).

**C. Vulnerability of xylem and outside-xylem tissue as a function of water potential (Fig. 4C,D in main text).** We also modeled the axial (along z) water potential gradient in the leaf using the hydraulic circuit as shown in Fig. 4C (main text). At a given location $z$ on the leaf, we consider the following pathway and resistances to water flux: axial flow, $J(z)$, through xylem experiences xylem resistance, $R_{\mathrm{xyl}}(\psi_{\mathrm{th}}^{\mathrm{xyl}})$, a function of xylem water potential, $\psi_{\mathrm{th}}^{\mathrm{xyl}}$; and the local evaporative flux (passing from xylem into mesophyll), $E$, experiences a resistance, $R_{\mathrm{ox}}(\psi_{\mathrm{th}}^{\mathrm{ox}})$, due to passage through bundle sheath cells and mesophyll; we assumed this resistance depends on the outside-xylem water potential($\psi_{\mathrm{th}}^{\mathrm{ox}}$). Mathematically, the governing equation for $\psi_{\mathrm{th}}^{\mathrm{ox}}$ is:

$$\frac{\partial \psi_{\mathrm{th}}^{\mathrm{ox}}}{\partial z} = -\frac{R_{\mathrm{xyl}}(\psi_{\mathrm{th}}^{\mathrm{xyl}})}{wL^2} J(z) - \frac{E}{L^2} R_{\mathrm{ox}}(\psi_{\mathrm{th}}^{\mathrm{ox}})\, dz, \tag{S19}$$

and the governing equation for $\psi_{\mathrm{th}}^{\mathrm{xyl}}$ is given by Eq. (S16). We take the axial flow, $J(z)$, through the leaf xylem to be governed by Eq. (S15). On substituting Eq. (S15) into Eq. (S19) and differentiating with respect to z, we obtain the equation for the evolution of water potential in terms of resistances and transpiration rate as a second order partial differential equation:

$$\frac{\partial^2 \psi_{\mathrm{th}}^{\mathrm{ox}}}{\partial z^2} = \frac{E}{L^2}(R_{\mathrm{xyl}}(\psi_{\mathrm{th}}^{\mathrm{xyl}}) - R_{\mathrm{ox}}(\psi_{\mathrm{th}}^{\mathrm{ox}})) - \frac{E}{L}\left(1 - \frac{z}{L}\right)\frac{\partial R_{\mathrm{xyl}}(\psi_{\mathrm{th}}^{\mathrm{xyl}})}{\partial z}. \tag{S20}$$

We used the following functional form for the resistances: (1) we fit a logistic function (Eq. (S17)) to the xylem vulnerability curve proposed by Li et al. ((25), shown in Fig. S15A). (2) We chose resistance from outside xylem, $R_{\mathrm{ox}}(\psi)$[ m$^2$.s.MPa/kg] to be a logistic function of $\psi$, such that,

$$R_{\mathrm{ox}}(\psi) = R_{\mathrm{ox}}^{\min}\left(1 + \left(\frac{\psi_{\mathrm{th}}^{\mathrm{ox}}}{\psi_{\mathrm{th},50\%}^{\mathrm{ox}}}\right)^a\right) \tag{S21}$$

where $R_{\mathrm{ox}}^{\min}$ is the minimum outside-xylem resistance (i.e., when $\psi_{\mathrm{th}}^{\mathrm{ox}} = 0$), and at $\psi_{\mathrm{th}}^{\mathrm{ox}} = \psi_{\mathrm{th},50\%}^{\mathrm{ox}}$, $R_{\mathrm{ox}}$ is twice that of $R_{\mathrm{ox}}^{\min}$.

We found a combination of $R_{\text{ox}}^{\text{min}}$ and $\psi_{\text{th},50\%}^{\text{ox}}$ such that the difference between $\psi_{\text{AQD}}^{\text{leaf}}$ and $\psi_{\text{th}}^{\text{ox}}$, such that:

$$\sum_{\text{Day }1,2,3} (\psi_{\text{AQD}}^{\text{leaf,mid}} - \psi_{\text{th}}^{\text{xyl}}(z = L/2)) + (\psi_{\text{AQD}}^{\text{leaf,tip}} - \psi_{\text{th}}^{\text{xyl}}(z = 5L/6)) \qquad \text{[S22]}$$

was minimized for both WW and WL plants for the range of transpiration rate measured experimentally. The minimization was performed by using the 'fminsearch' solver in MATLAB software. Correspondingly, we obtained range for $R_{\text{ox}}$ for the range of measured $E$, shown as shaded blue region in Fig. S15A. Upon substituting substituting Eq. (S17) and Eq. (S21), we solved coupled differential equations, Eq. (S16) and Eq. (S20) numerically to obtain $\psi_{\text{th}}^{\text{xyl}}$ and $\psi_{\text{th}}^{\text{ox}}$ as a function of $z$ for the case of well-watered (WW) and water-limited (WL) plants and plot it as shown in Fig. 4D, with the following boundary conditions as derived from the hydraulic architecture shown in Fig. 4C (main text): (1) at $z = L/6$, $\psi_{\text{th}}^{\text{xyl}} = \psi_{\text{AQD}}^{\text{Node}} + E \times R_{\text{ox}}(\psi_{\text{AQD}}^{\text{Node}})$ (2) at $z = L/6$, $\psi_{\text{th}}^{\text{ox}} = \psi_{\text{AQD}}^{\text{Node}}$, and (3) at $z = L/6$, we make a finite difference approximation for the second term on the right side of the Eq. (S19) such that:

$$\left.\frac{\partial \psi_{\text{ox}}}{\partial z}\right|_{z=L/6} = -\left.\frac{R_{\text{xyl}}(\psi)}{wL^2} J(z)\right|_{z=L/6,\, \psi=\psi_{\text{th}}^{\text{xyl}}} - \left.\frac{E}{L^2} R_{\text{ox}}(\psi)\, \Delta z\right|_{\Delta z = L/6,\, \psi=\psi_{\text{AQD}}^{\text{Node}}}. \qquad \text{[S23]}$$

The corresponding prediction of $\psi_{\text{th}}^{\text{ox}}$ is in agreement with the water potential measured using AquaDust at node, mid and tip of the maize leaf (Fig. 4D, main text). The obtained parameter values of $\psi$-dependence for extra-vascular resistance are similar to the values reported for the mesophyll resistance obtained for other species based on the vacuum pressure method and modeling studies (27, 28): briefly, $\psi_{\text{th},50\%}^{\text{ox}} < \psi_{\text{th},50\%}^{\text{xyl}}$ and $R_{\text{ox}}$ accounts for $75 - 100\%$ of the leaf resistance under mild water stress.

**D. Prediction of diurnal water potential corresponding to Fig. 5 (main text).** We used AquaDust to measure the diurnal variation in leaf water potential and estimated transpiration rate ($E$) using eddy flux measurements of net water vapor flux above the canopy in a well-irrigated plot at Cornell Musgrave Research Farm (Location: 42°43′N, 76°39′W). Here, we describe the calculation of theoretical prediction for leaf water potential using the hydraulic architecture for the maize plant as shown in Fig. S16. The measurements reported in Fig. 5B (main text) were performed at the tip of the leaves and, as before, we assume that they report the water potential in the mesophyll, outside the xylem, as indicated by the red point on the diagram in Fig. S16.

We take the soil to be saturated ($\psi^{\text{soil}} = 0$), and we assume root and stem have negligible resistance relative to that in leaves. This second assumption is consistent with a study done in well-watered potted maize plants with a continuous measurement of stem water potential using a micro-tensiometer (29). We note that some studies in the literature have inferred a more equal partitioning of resistance between the roots and soil and the stem and leaves (30, 31), although we are unaware of direct, in situ measurements that verify the actual partitioning.

We used the values of xylem resistance ($R_{\mathrm{xyl}}(\psi_{\mathrm{th}}^{\mathrm{xyl}})$) and outside-xylem resistance ($R_{\mathrm{ox}}(\psi_{\mathrm{th}}^{\mathrm{ox}})$) based on our inference from the gradient of water potential along the leaf (Sec. S5C, Fig. S15A). We reiterate on those briefly here: (1) $R_{\mathrm{xyl}}(\psi_{\mathrm{th}}^{\mathrm{xyl}})$ is obtained by fitting a logistic function to the xylem vulnerability curve proposed by Li et al. ((25), shown in Fig. S15). (2) We used the inferred mean value of outside-xylem resistance (noted in the table and shown as a solid blue line, Fig. S15A) to generate the mean theoretical prediction for leaf water potential (shown as solid blue dots in Fig. 5 - main text); we used the minimum and maximum values of outside xylem resistance (noted in the table and shown as shaded blue region, Fig. S15A) to generate the range for theoretical prediction of $\psi_{\mathrm{th}}^{\mathrm{ox}}$ (shown as shaded blue region in Fig. 5B - main text).

We solve for diurnal evolution of $\psi_{\mathrm{th}}^{\mathrm{xyl}}$ and $\psi_{\mathrm{th}}^{\mathrm{ox}}$ at the tip of the leaf with hourly-averaged measurements of $E$ (Fig. 5A - main text) using coupled differential equations: Eq. (S16) and Eq. (S20), with the following boundary conditions at $z = 0$: (1) $\psi_{\mathrm{th}}^{\mathrm{xyl}} = \psi^{\mathrm{soil}} \equiv 0$; (2) $\psi_{\mathrm{th}}^{\mathrm{ox}} = \psi_{\mathrm{th}}^{\mathrm{xyl}} - E \times R_{\mathrm{ox}}(\psi_{\mathrm{th}}^{\mathrm{ox}})$; (3) first order derivative for $\psi_{\mathrm{th}}^{\mathrm{ox}}$ with respect to $z$ given by Eq. (S23).

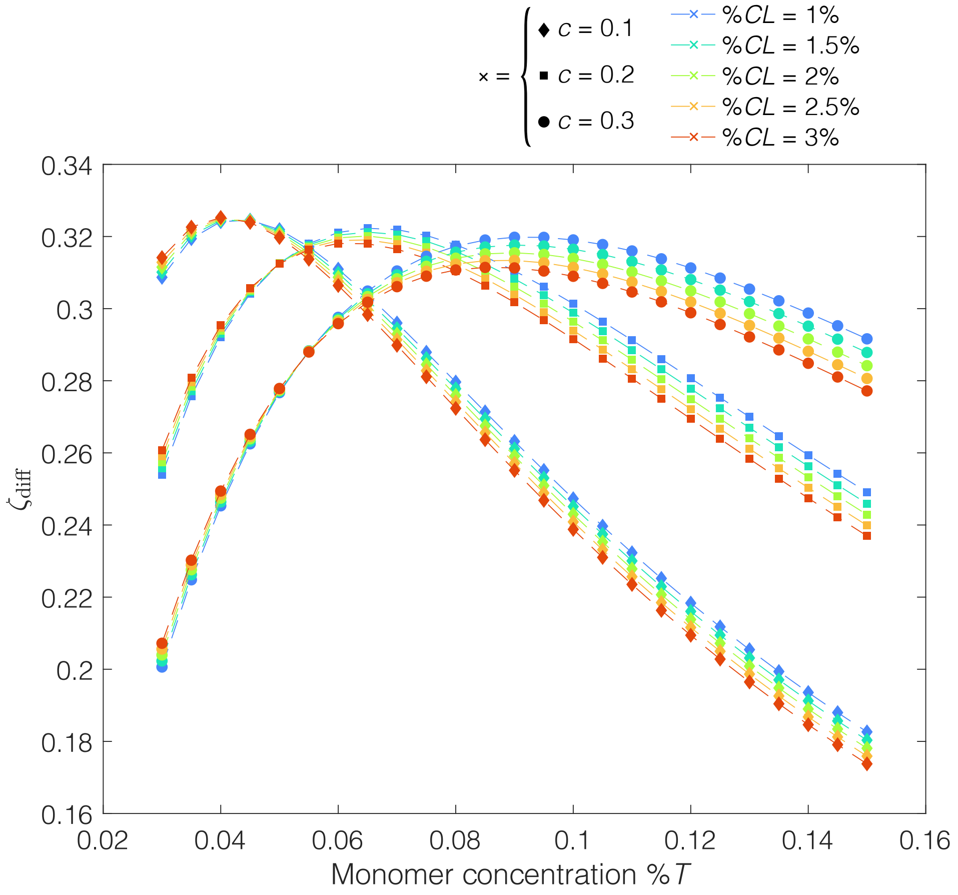

**Fig. S1.** Theoretical model for determining monomer and cross-linker concentration to maximize AquaDust sensitivity: $\zeta_{\mathrm{diff}}$ (Eq. (S12)) is plotted against percentage monomer concentration, $\%T$, for different values of relative cross-linker concentration, $\%CL$, and fit parameter, $c$.

 

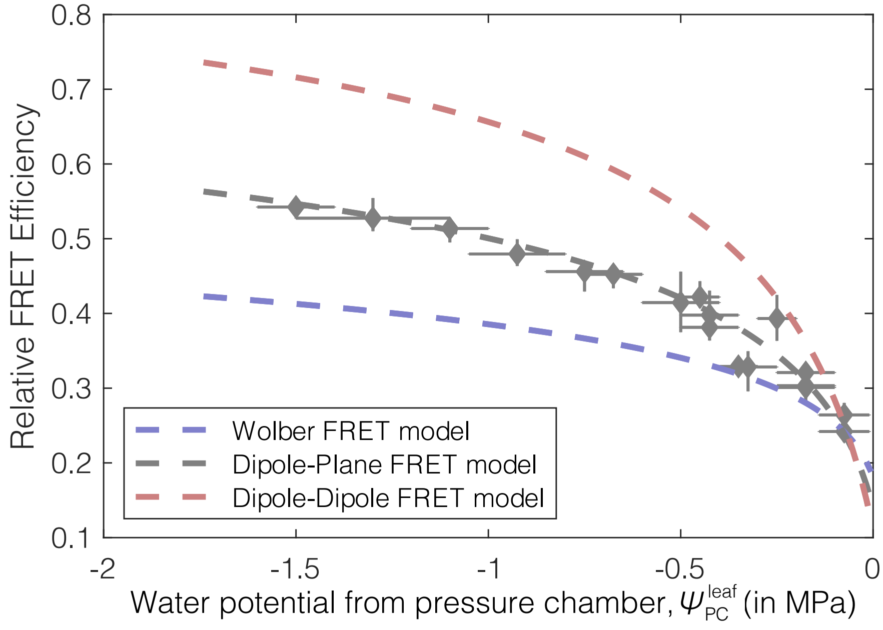

**Fig. S2.** Comparison of different models for FRET prediction: Experimental relative FRET Efficiency ($\zeta_{\mathrm{exp}}$) as a function of $\psi_{\mathrm{PC}}^{\mathrm{leaf}}$ (as shown in Fig. 3C - main text) is plotted as grey diamonds. The vertical error bars represent the range of $\zeta_{\mathrm{exp}}$ from AquaDust and the horizontal error bars represent range of water potential from pressure chamber (see Sec. S4 I, Table S3). The theoretical prediction, $\zeta_{\mathrm{th}}$ from three different models: Wolber FRET Model, Dipole-Plane FRET Model and Dipole-Dipole FRET model is plotted as shown for comparison. The fit parameter in each of these models is obtained by forcing $\zeta_{\mathrm{th}} = \zeta_{\mathrm{exp}}$ to the experimental data closest to saturation (here, $\psi_{\mathrm{PC}}^{\mathrm{leaf}} = -0.08$ MPa).

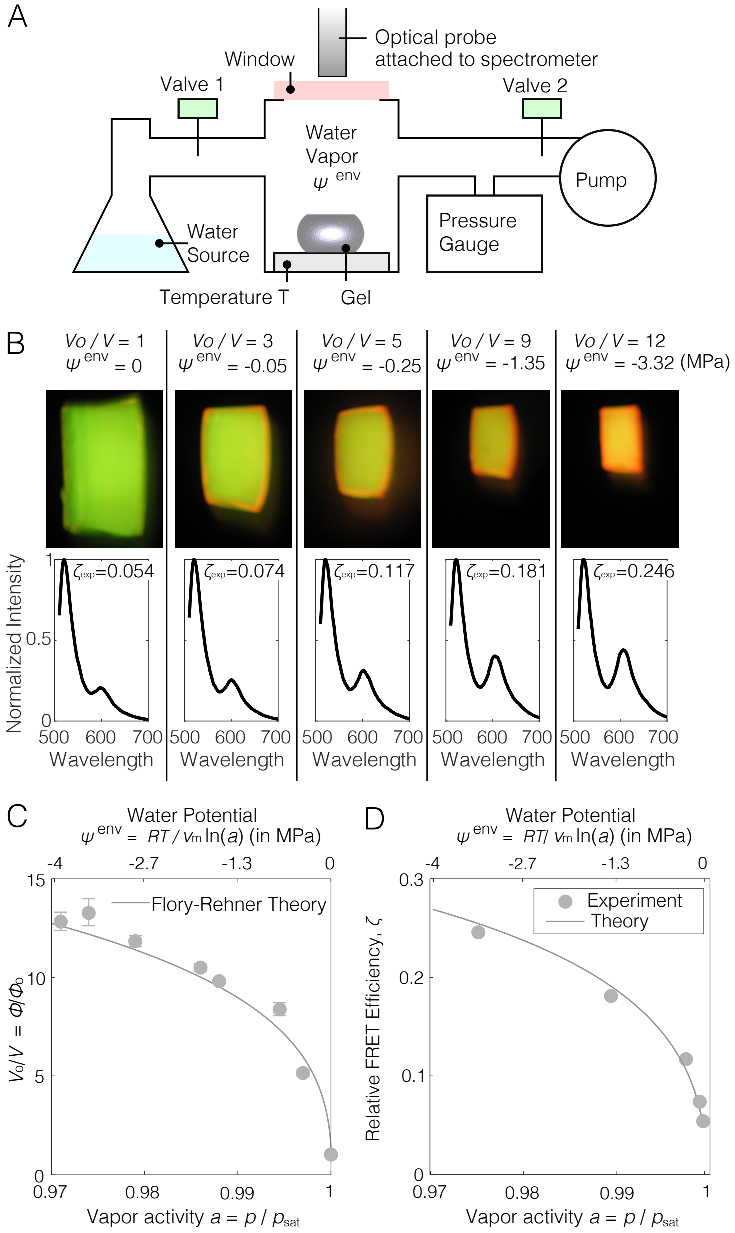

**Fig. S3.** Characterization of bulk swelling and fluorescence of gel matrix using vapor pressure to define $\psi^{\rm env}$. (A) Setup used to measure gel response as function of imposed water potential, $\psi^{\rm env}$. See (17) for detailed description of the setup. (B) Demonstration of water potential sensing mechanism using a bulk gel undergoing change in volume (Volume at $\psi^{\rm env} = 0$: $V_{\rm o}$, Volume when $\psi^{\rm env} < 0$: $V$) and color upon changing the $\psi^{\rm env}$. The bottom panel shows the emission spectra from the bulk gels corresponding to the images. As water potential decreases, relative intensity of the acceptor dye ($\sim 605$ nm) increases. (C) Change in the volume of bulk gel with change in vapor activity ($a$, bottom axis) and water potential ($\psi^{\rm env}$, top axis) as compared with theoretical prediction (Flory Rehner model, Eq. (S1)). $V_o$ is the initial volume of gel and $V$ is the final volume of gel for a given vapor activity; (D) Experimental FRET Efficiency ($\zeta_{\rm exp}$) from bulk gels as compared to that of the theoretical prediction ($\zeta_{\rm th}$), (Eq. (S10)). Parameter values for fits in (C) and (D) are provided in Table S1.

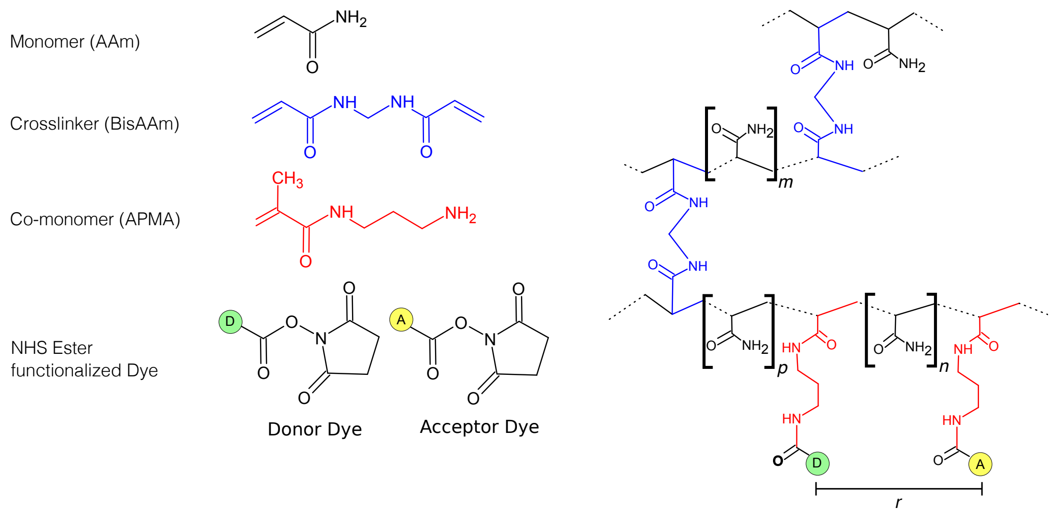

**Fig. S4.** Chemical composition of gel matrix: Acrylamide polymer chain is interspersed with N-3 Aminopropyl methacrylamide (APMA) which is conjugated to donor and acceptor fluorophores, shown in green (either AF488 for macroscopic gel or OG for AquaDust-nanogel synthesis (not shown)) and yellow (either AF568 for macroscopic gel RH for AquaDust-nanogel synthesis (not shown)) circles modified with N-hydroxy succinimidyl ester.

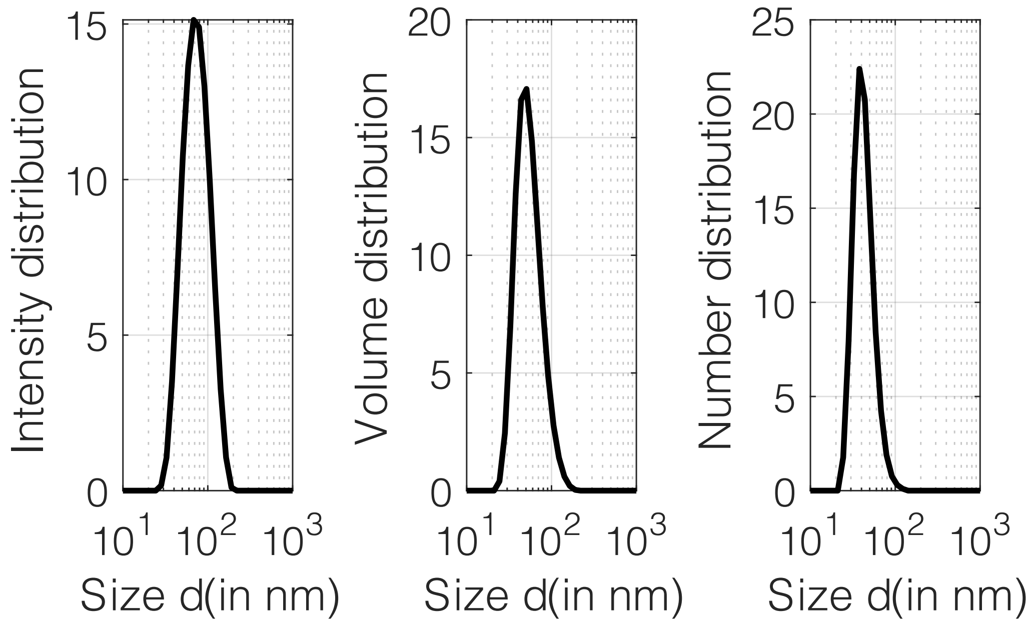

**Fig. S5.** AquaDust size distribution: Size (diameter) distribution with respect to intensity of scattering in dynamic light scattering measurement, with respect to the number of particles, and with respect to the volume of particles averaged over three measurements.

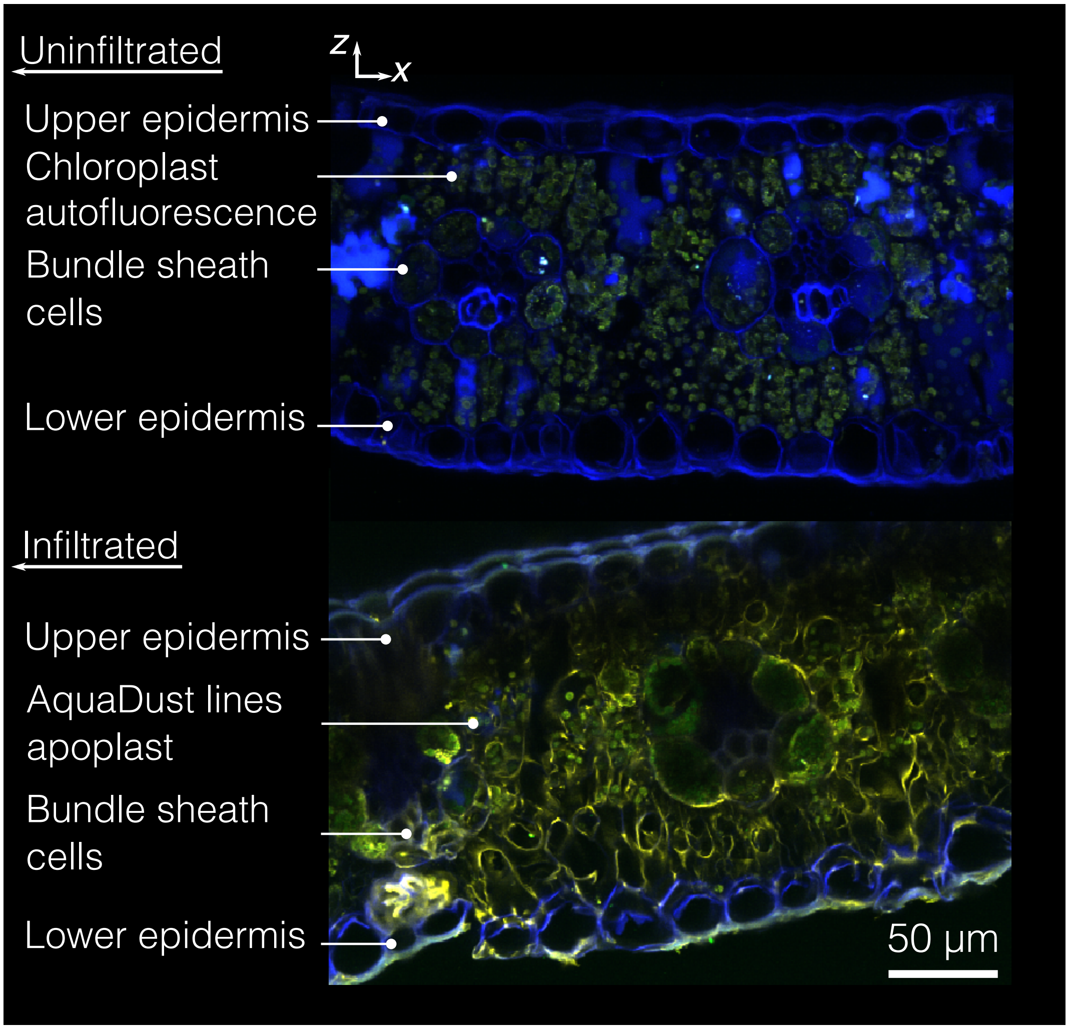

**Fig. S6.** Cross-section view of AquaDust distribution within mesophyll: Cross-section of uninfiltrated and infiltrated zone were imaged under confocal microscope where AquaDust adds to the fluorescence in falsely-colored yellow channel (Scale bar: 50 $\mu$m).

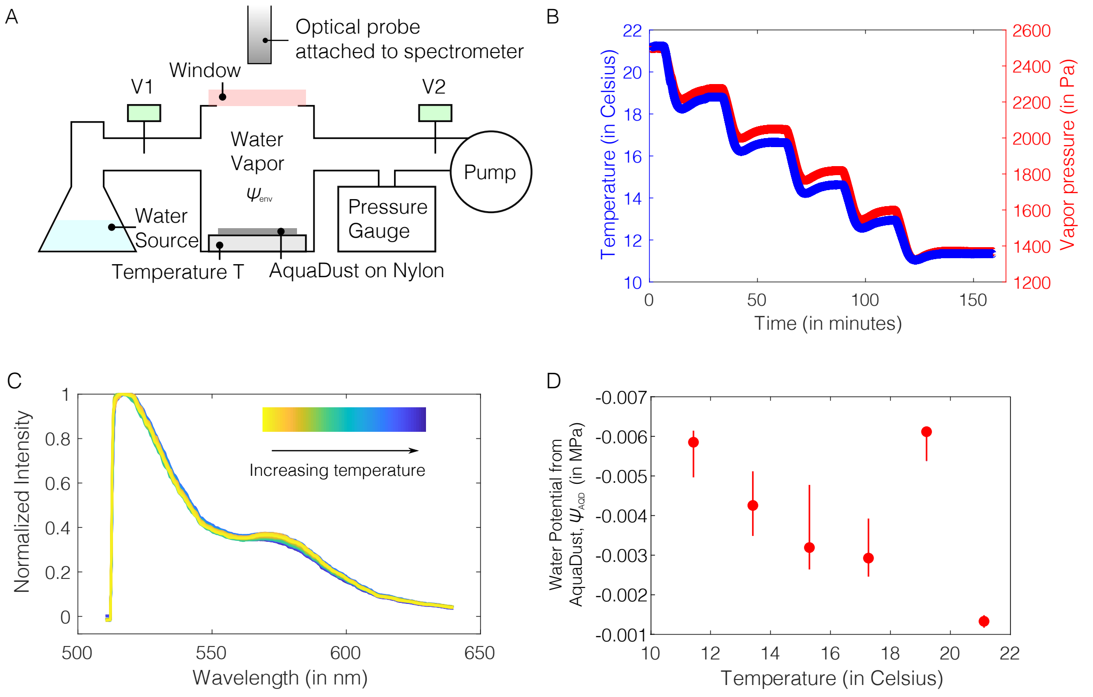

**Fig. S7.** AquaDust response to temperature: (A) Setup used to measure AquaDust response as function of temperature. (B) Step changes made in temperature, $T$, and vapor pressure such that the vapor pressure ($p$) equals the saturation vapor pressure ($p_{\mathrm{sat}}$) ($\psi^{\mathrm{env}} = 0$) at given temperature $T$ is plotted as a function of time and the emission spectra (C) was recorded 30 seconds before step change in temperature. (D) Value of water potential extracted from spectra in (C) based on calibration performed at 24°C .

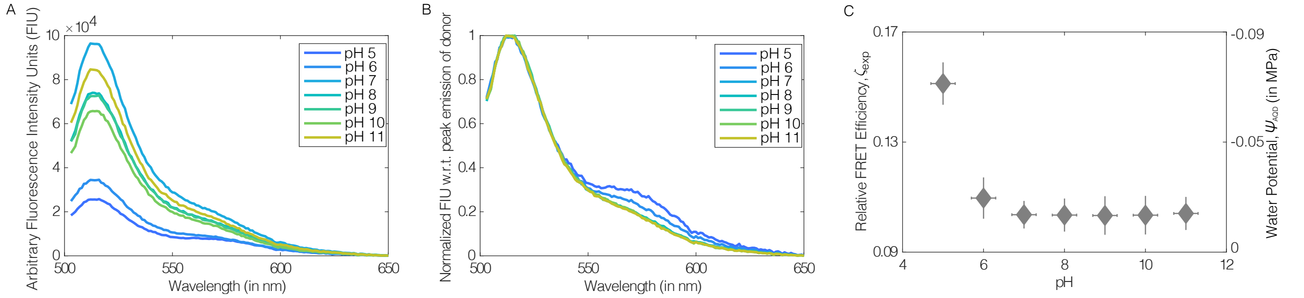

**Fig. S8.** AquaDust response to pH: (A) Raw fluorescence spectra and, (B) fluorescence spectra normalized with respect to peak emission intensity, as measured from AquaDust as a function of pH of buffer solutions. (C) Value of experimental relative FRET Efficiency and corresponding water potential as extracted from spectra in (B) based on calibration performed at pH 7.

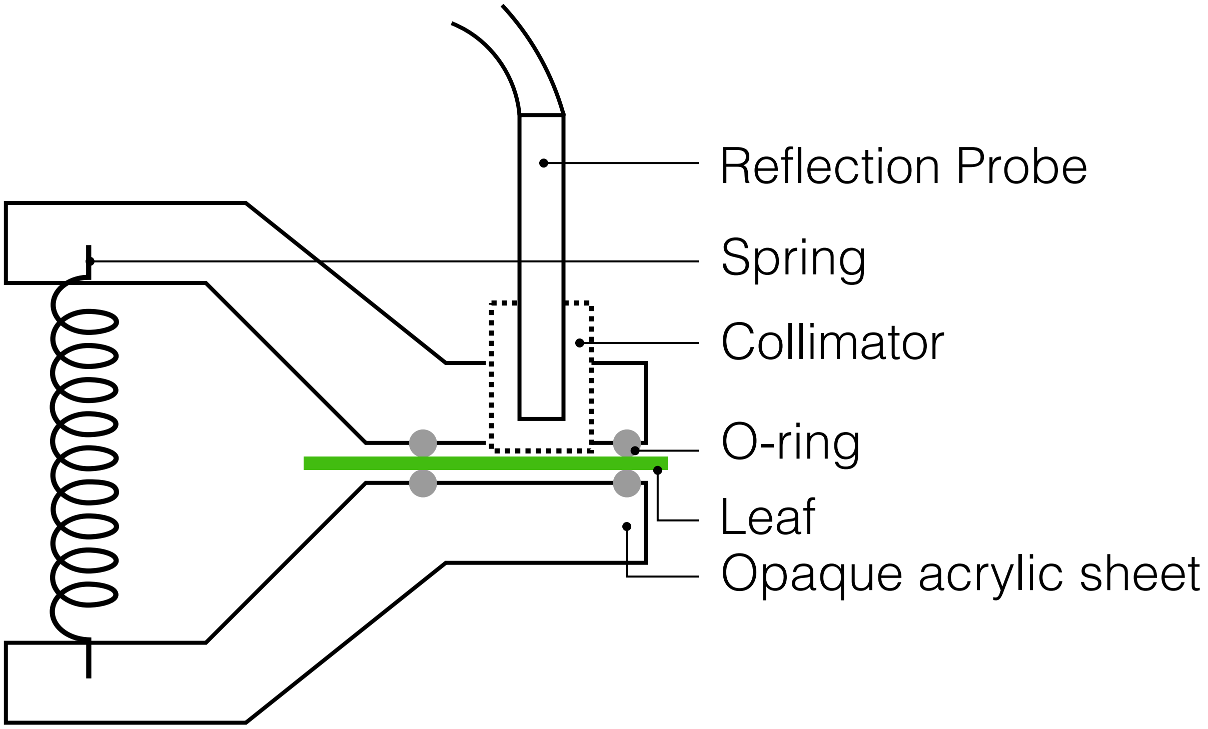

**Fig. S9.** Leaf clamp designed to position the reflection probe to excite and collect AquaDust fluorescence, O-rings separate the contact of fiber bundle with the site of interrogation (See details in Sec. S4H).

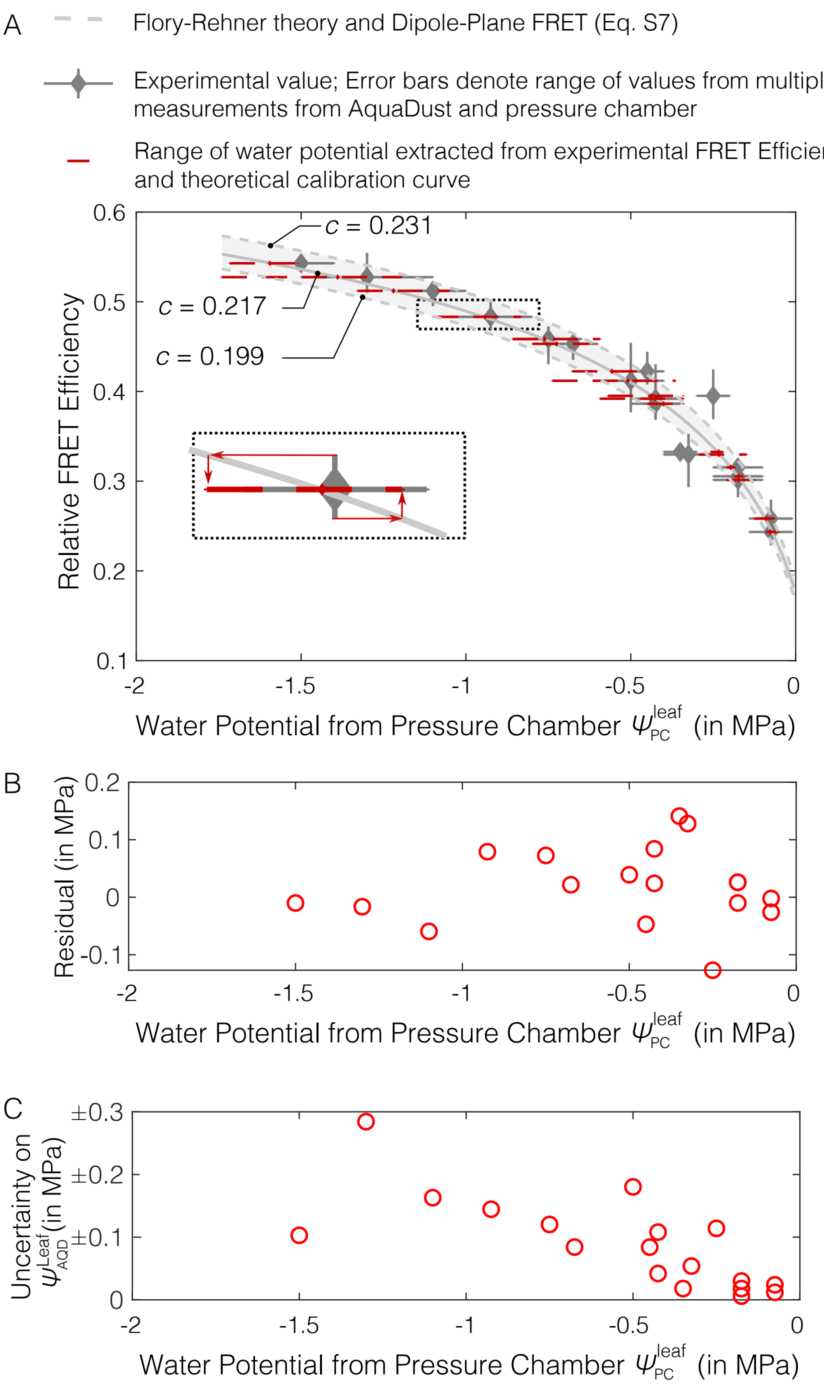

**Fig. S10.** Accuracy of measurements from AquaDust: (A) Experimental relative FRET Efficiency ($\zeta_{\mathrm{exp}}$) as a function of $\psi_{\mathrm{PC}}^{\mathrm{leaf}}$ (as shown in Figure 3C) is plotted as grey diamonds. Bounds on the theoretical prediction obtained on the fitting parameter, $c$; dotted gray lines correspond to the $c = 0.199$ and $c = 0.231$, calculated upon minimizing the difference between minimum/maximum $\psi_{\mathrm{AQD}}^{\mathrm{leaf}}$ and the water potential from pressure chamber $\psi_{\mathrm{PC}}$ respectively, solid gray line correspond to $c = 0.217$ as calculated upon minimizing the difference between mean experimental values of water potential from AquaDust and the water potential from pressure chamber. (B) Difference between mean value of water potential as extracted from the mean value of FRET Efficiency ($\psi_{\mathrm{AQD}}^{\mathrm{leaf}}$) and the mean value of water potential from pressure chamber ($\psi_{\mathrm{PC}}^{\mathrm{leaf}}$). (C) Experimental range of water potential ($\psi_{\mathrm{AQD}}^{\mathrm{leaf}}$) is obtained from the range of AquaDust FRET Efficiency using the theoretical prediction curve as shown in (A, red-dashed lines), and is plotted as uncertainty.

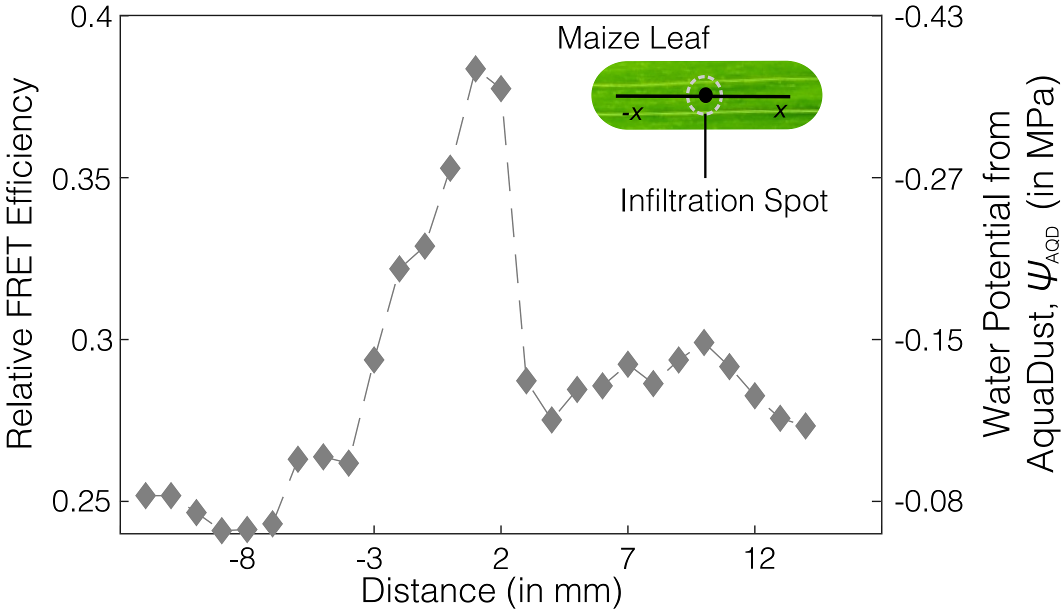

**Fig. S11.** Effect of cuticle damage during infiltration on water potential measured using AquaDust: FRET Efficiency and corresponding water potential based on calibration (Fig. 3C - main text) is plotted as function of distance from spot of infiltration (Maximum distance affected by the damage due to infiltration $\sim \pm$ 3 mm).

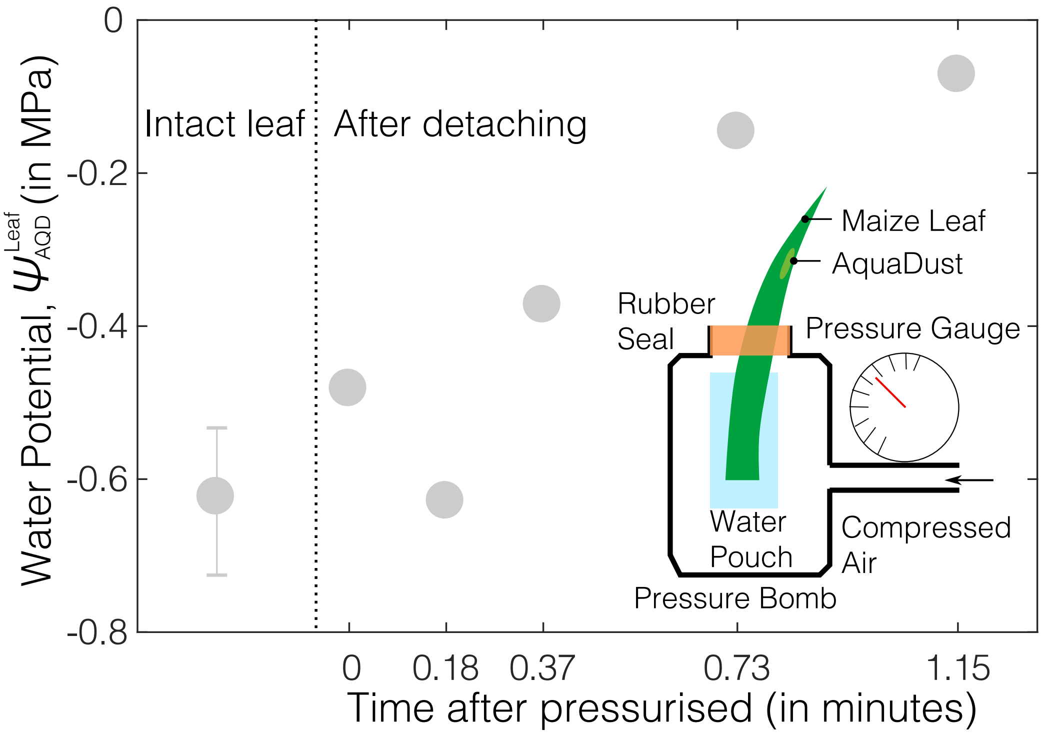

**Fig. S12.** Temporal response of AquaDust in leaves: Response time of AquaDust is measured by placing cut-out maize leaf with exposed region in water pouch and pressuring it at +0.2 MPa in a pressure chamber, water potential from AquaDust is plotted against time after it was pressurized.

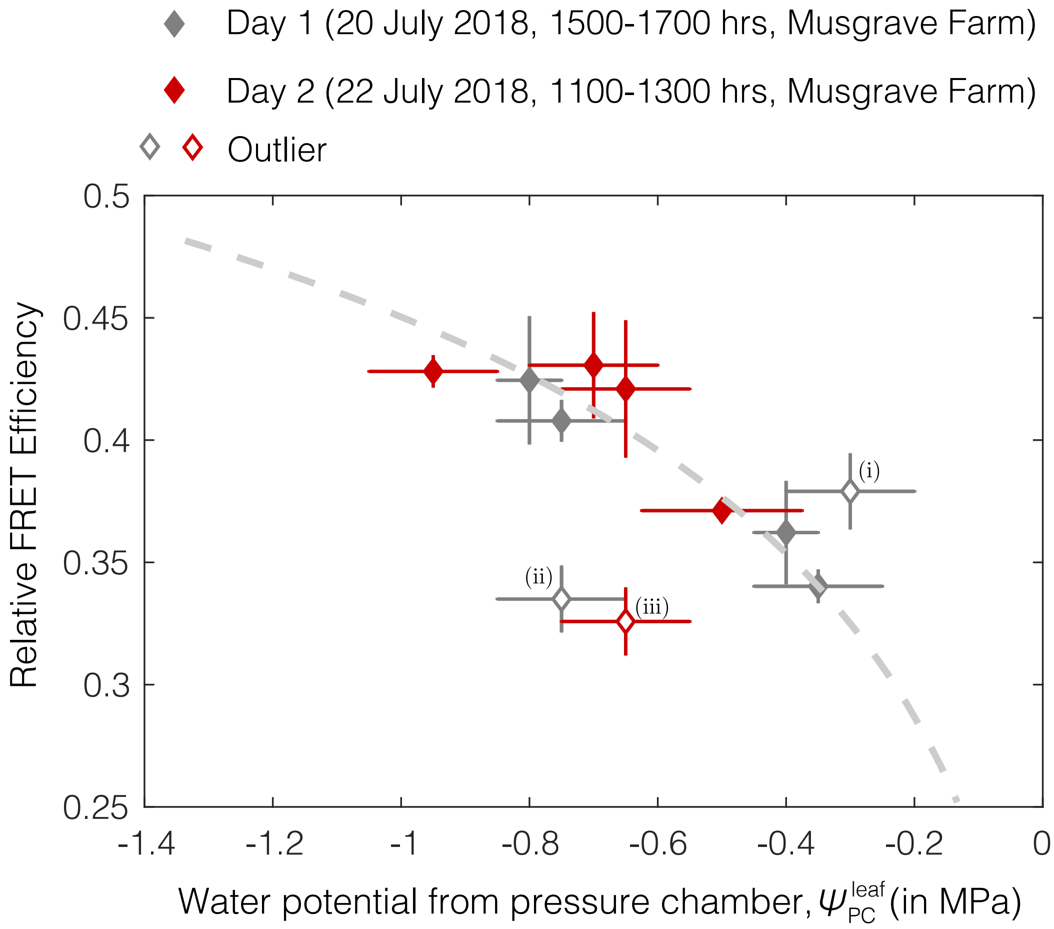

**Fig. S13.** In-field comparison of water potential from AquaDust and Scholander pressure chamber: Relative FRET Efficiency from AquaDust is plotted against water potential from pressure chamber, $\psi_{\mathrm{PC}}^{\mathrm{leaf}}$. Theoretical prediction (dashed line) as obtained from the Flory-Rehner theory and Dipole-Plane FRET model is plotted against water potential.

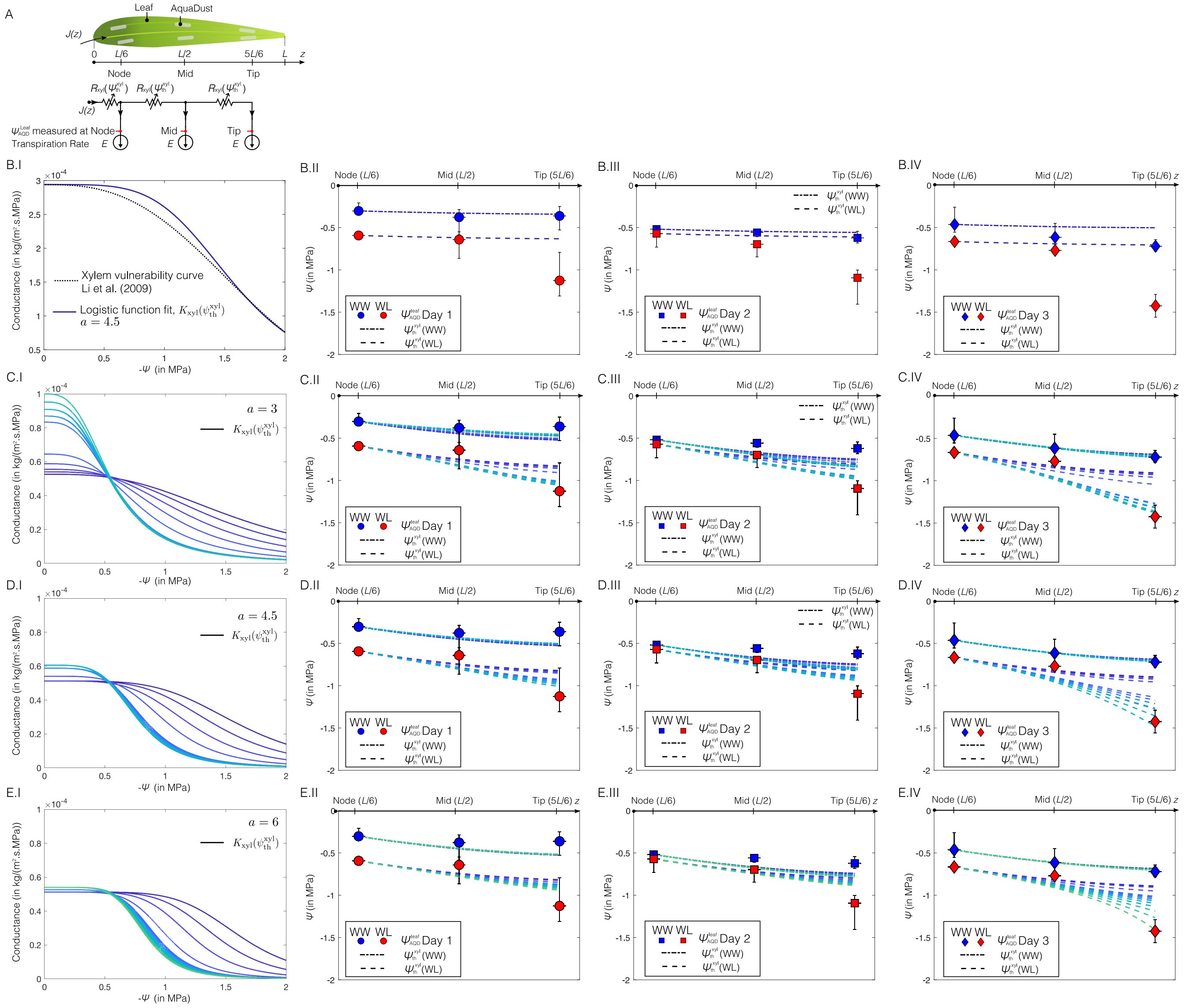

**Fig. S14.** Model for water potential gradients in leaf considering only xylem resistance (see Sec. S5B): (A) Resistance-based model to predict the water potential as a function of xylem resistance ($R_{\mathrm{xyl}}$) to axial flux $J(z)$ with constant transpiration flux, $E$, where water potential from AquaDust, $\psi_{\mathrm{AQD}}^{\mathrm{Leaf}}$, is measured at the tip, mid and node of the leaf. (B) Prediction of xylem water potential ($\psi_{\mathrm{th}}^{\mathrm{xyl}}$) (Eq. (S16)) with a xylem conductance, $K_{\mathrm{xyl}}(\psi_{\mathrm{th}}^{\mathrm{xyl}}) = 1/R_{\mathrm{xyl}}(\psi_{\mathrm{th}}^{\mathrm{xyl}})$ given by logistic function (Eq. (S17)) fit to the vulnerability curve for xylem obtained by Li et al. (25) (B.I) and comparison with $\psi_{\mathrm{AQD}}^{\mathrm{leaf}}$ measured at node, mid, and tip of the leaf with maize plant in well-watered (WW) condition and for plants left unwatered (water-limited, WL) for (B.II) 1 day (Day 1), (B.III) 2 days (Day 2) and (B.IV) 3 days (Day 3). $K_{\mathrm{xyl}}(\psi_{\mathrm{th}}^{\mathrm{xyl}}) = 1/R_{\mathrm{xyl}}(\psi_{\mathrm{th}}^{\mathrm{xyl}})$ as a function of $\psi_{\mathrm{th}}^{\mathrm{xyl}}$ was calculated such that the difference between $\psi_{\mathrm{th}}^{\mathrm{xyl}}$ (Eq. (S16)) and $\psi_{\mathrm{AQD}}^{\mathrm{leaf}}$ was minimum for WW case with different logistic growth factors (C.I) $a = 3$, (D.I) $a = 4.5$, and (E.I) $a = 6$; and respective prediction (lines of same color) of $\psi_{\mathrm{th}}^{\mathrm{xyl}}$ was compared with $\psi_{\mathrm{AQD}}^{\mathrm{leaf}}$ measured for WW and WL plants on Day 1 (C.II, D.II, E.II), Day 2 (C.III, D.III, E.III) and Day 3 (C.IV, D.IV, E.IV).

Xylem and outside-xylem resistance:

$$R\,(\Psi_{th}) = R^{min}\left(1+\frac{\Psi_{th}}{\Psi_{th,50\%}}\right)^{a}$$

| | $ET$ in kg/(m$^2$.s) | $R^{min}$ in (m$^2$.s.MPa)/kg | $\Psi_{th,50\%}$ (in MPa) | $a$ |
|---|---|---|---|---|
| Xylem (xyl) | | $3.47\times10^{3}$ | -1.58 | 4.5 |
| Outside-Xylem Resistance (ox) | $3.4\times10^{-5}$(Min) | $4.4\times10^{3}$ | -0.44 | 4.5 |
| | $4.2\times10^{-5}$(Mean) | $3.7\times10^{3}$ | -0.45 | 4.5 |
| | $5.1\times10^{-5}$(Max) | $3.2\times10^{3}$ | -0.46 | 4.5 |

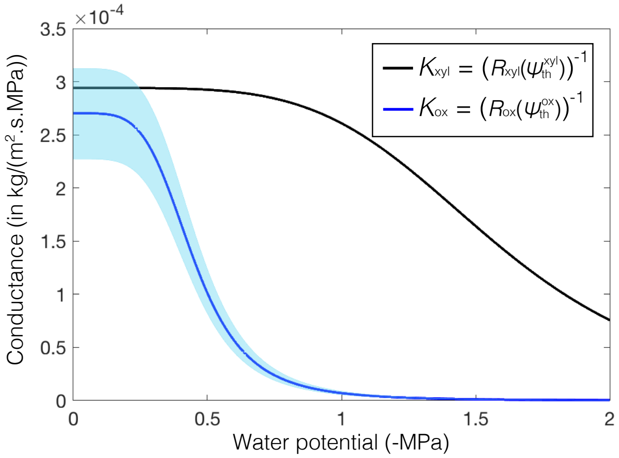

**Fig. S15.** Mesophyll resistance and corresponding prediction of outside-xylem water potential: Logistic function fit for xylem conductance, $K_{\mathrm{xyl}} = 1/R_{\mathrm{xyl}}(\psi_{\mathrm{th}}^{\mathrm{xyl}})$ to the vulnerability curve for xylem obtained by Li et al. (25) and outside-xylem conductance ($K_{\mathrm{ox}} = 1/R_{\mathrm{ox}}(\psi_{\mathrm{th}}^{\mathrm{ox}})$) calculated such that the difference between $\psi_{\mathrm{th}}^{\mathrm{ox}}$ (Eq. (S16)) and $\psi_{\mathrm{AQD}}^{\mathrm{leaf}}$ was minimum for both WW and WL plants, with solid line corresponding to the mean transpiration rate $\left(4.2\times10^{-5}\ \mathrm{kg}/(\mathrm{m}^2.\mathrm{s})\right)$ and the upper and lower bound of the shaded region as calculated corresponding to the minimum and maximum values of transpiration rate ($3.4\times10^{-5}\ \mathrm{kg}/(\mathrm{m}^2.\mathrm{s})$ and $5.1\times10^{-5}\ \mathrm{kg}/(\mathrm{m}^2.\mathrm{s})$ respectively) as reported in the table.

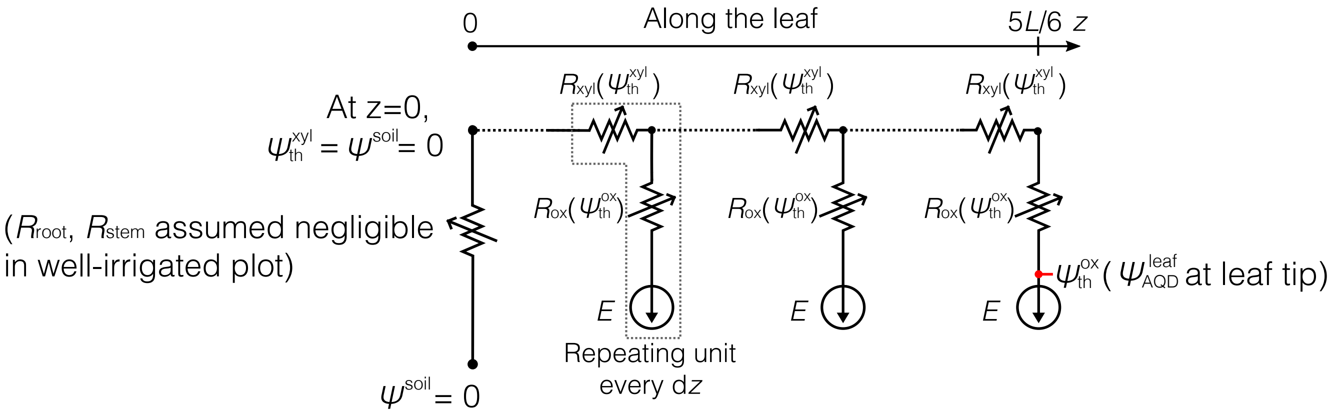

**Fig. S16.** Resistance-based hydraulic model for a well-watered maize plant used to calculate leaf water potential based on measured transpiration ($E$, Fig. 5A - main text), assuming negligible root and stem resistances and values of xylem resistance ($R_{\rm xyl}$) and outside-xylem resistance ($R_{\rm ox}$) inferred from gradients of water potential in transpiring leaf (Fig. S15).

### Table S1. AquaDust gel composition and parameters in Eq. (S1)

| | Name (Abbreviation/Symbol) | Composition/Value |
|---|---|---|
| 1. Monomer concentration | Acrylamide (AAm) | $w_{\mathrm{mon1}} = 0.055[\mathrm{gm/cm^3}]$ |
| 2. Co-monomer concentration | N-aminopropyl methacrylamide (APMA) | $w_{\mathrm{mon2}} = 0.006[\mathrm{gm/cm^3}]$ |
| 3. Total monomer concentration | $w_{\mathrm{mon}} = w_{\mathrm{mon1}} + w_{\mathrm{mon2}}$ | $w_{\mathrm{mon}} = 0.061[\mathrm{gm/cm^3}]$ |
| 4. Cross-linker concentration | N,N-methylenebisacrylamide (BisAAm) | $w_{\mathrm{cl}} = 0.0018[\mathrm{gm/cm^3}]$ |
| 5. Theta temperature | $\theta$ | $\theta = 275[K]$ |
| 6. Gel temperature | $T$ | $T = 288[K]$ |
| 7. Molar weight of monomer | $mw_{\mathrm{mon1}}$ | $mw_{\mathrm{mon1}} = 71.08[\mathrm{gm/mol}]$ |
| 8. Molar weight of co-monomer | $mw_{\mathrm{mon2}}$ | $mw_{\mathrm{mon2}} = 178.66[\mathrm{gm/mol}]$ |
| 9. Specific volume of monomer | $v_{\mathrm{mon1}}$ | $v_{\mathrm{mon1}} = 0.756[\mathrm{cm^3/gm}]$ |
| 10. Specific volume of co-monomer | $v_{\mathrm{mon2}}$ | $v_{\mathrm{mon2}} = 0.812[\mathrm{cm^3/gm}]$ |
| 11. Molar weight of crosslinker | $mw_{\mathrm{cl}}$ | $mw_{\mathrm{cl}} = 154.17[\mathrm{gm/mol}]$ |
| 12. Specific volume of crosslinker | $v_{\mathrm{cl}}$ | $v_{\mathrm{cl}} = 0.8097[\mathrm{cm^3/gm}]$ |
| 13. Molar weight of water | $mw_{\mathrm{w}}$ | $mw_{\mathrm{w}} = 18.0153[\mathrm{gm/mol}]$ |
| 14. Specific volume of water | $v_{\mathrm{w}}$ | $v_{\mathrm{w}} = 1.001[\mathrm{cm^3/gm}]$ |

### Table S2. Effect of AquaDust infiltration on the physiological parameters of the maize leaf

| Experiment No | Plant No | Leaf No | $G_{H_2O}$ (mmol/(m$^2$.s)) | | A ($\mu$mol/(m$^2$.s)) | | E (mmol/(m$^2$.s)) | |
|---|---|---|---|---|---|---|---|---|
| | | | AquaDust | Control | AquaDust | Control | AquaDust | Control |
| 1 | 1 | 7 | 118.2 | 129.8 | 13.9 | 13.4 | 1.71 | 1.80 |
| | 2 | 7 | 121.8 | 120.9 | 15.0 | 15.0 | 1.80 | 1.76 |
| | 3 | 7 | 112.2 | 82.7 | 12.2 | 8.2 | 1.69 | 1.28 |
| | 4 | 7 | 170.1 | 168.6 | 16.0 | 15.5 | 2.46 | 2.40 |
| | 5 | 7 | 171.8 | 111.3 | 16.1 | 12.4 | 2.43 | 1.68 |
| Paired sample two-tailed t-test | | P-value | 0.281 | | 0.113 | | 0.207 | |

| Experiment No | Plant No | Leaf No | $G_{H_2O}$ (mmol/(m$^2$.s)) | | A ($\mu$mol/(m$^2$.s)) | | E (mmol/(m$^2$.s)) | |
|---|---|---|---|---|---|---|---|---|
| | | | AquaDust | Water | AquaDust | Water | AquaDust | Water |
| 2 | 5 | 6 | 123.9 | 188.7 | 11.5 | 16.0 | 1.87 | 2.65 |
| | 1 | 6 | 126.3 | 121.8 | 12.1 | 12.4 | 1.88 | 1.82 |
| | 2 | 6 | 116.5 | 111.3 | 11.8 | 11.2 | 1.73 | 1.75 |
| | 4 | 6 | 98.7 | 89.3 | 12.8 | 10.3 | 1.52 | 1.40 |
| Paired sample two-tailed t-test | | P-value | 0.567 | | 0.791 | | 0.514 | |

### Table S3. Numerical values of data shown in Fig. 3C

| $\psi_{\mathrm{PC}}^{\mathrm{leaf}}$ (in MPa) | | | FRET Efficiency $\zeta_{\mathrm{exp}}$ | | |
|---|---|---|---|---|---|
| Minimum | Mean | Maximum | Minimum | Mean | Maximum |
| -0.03 | -0.08 | -0.12 | 0.228 | 0.243 | 0.256 |
| -0.05 | -0.08 | -0.14 | 0.247 | 0.258 | 0.28 |
| -0.05 | -0.18 | -0.20 | 0.298 | 0.305 | 0.308 |
| -0.08 | -0.18 | -0.24 | 0.312 | 0.315 | 0.331 |
| -0.10 | -0.20 | -0.25 | 0.282 | 0.301 | 0.320 |
| -0.20 | -0.25 | -0.30 | 0.363 | 0.395 | 0.425 |
| -0.25 | -0.33 | -0.40 | 0.296 | 0.330 | 0.349 |
| -0.30 | -0.35 | -0.40 | 0.321 | 0.333 | 0.337 |
| -0.35 | -0.43 | -0.50 | 0.376 | 0.392 | 0.431 |
| -0.35 | -0.43 | -0.52 | 0.364 | 0.386 | 0.392 |
| -0.40 | -0.45 | -0.50 | 0.408 | 0.422 | 0.443 |
| -0.40 | -0.50 | -0.60 | 0.375 | 0.412 | 0.456 |
| -0.60 | -0.68 | -0.75 | 0.434 | 0.453 | 0.463 |
| -0.65 | -0.75 | -0.85 | 0.429 | 0.458 | 0.47 |
| -0.80 | -0.93 | -1.05 | 0.463 | 0.483 | 0.499 |
| -1.05 | -1.10 | -1.20 | 0.495 | 0.512 | 0.527 |
| -1.10 | -1.30 | -1.53 | 0.51 | 0.528 | 0.554 |
| -1.30 | -1.50 | -1.65 | 0.536 | 0.543 | 0.551 |

*Macromolecules* 13(4):977–983.

19. Clark HA, Hoyer M, Philbert MA, Kopelman R (1999) Optical nanosensors for chemical analysis inside single living cells. 1. Fabrication, characterization, and methods for intracellular delivery of PEBBLE sensors. *Analytical Chemistry* 71(21):4831–4836.
20. Clark HA, Kopelman R, Tjalkens R, Philbert MA (1999) Optical nanosensors for chemical analysis inside single living cells. 2. Sensors for pH and calcium and the intracellular application of PEBBLE sensors. *Analytical Chemistry* 71(21):4837–4843.
21. Okano T, Bae YH, Jacobs H, Kim SW (1990) Thermally on-off switching polymers for drug permeation and release. *Journal of Controlled Release* 11(1-3):255–265.
22. Owens DE, et al. (2007) Thermally responsive swelling properties of polyacrylamide/poly(acrylic acid) interpenetrating polymer network nanoparticles. *Macromolecules* 40(20):7306–7310.
23. Sun H, Almdal K, Andresen TL (2011) Expanding the dynamic measurement range for polymeric nanoparticle pH sensors. *Chemical Communications* 47(18):5268–5270.
24. Turner NC (1974) Stomatal Behavior and Water Status of Maize, Sorghum, and Tobacco under Field Conditions. *Plant Physiology* 53(3):360–365.
25. Li Y, Sperry JS, Shao M (2009) Hydraulic conductance and vulnerability to cavitation in corn (Zea mays L.) hybrids of differing drought resistance. *Frontiers in Plant Science* 66(2):341–346.
26. Chunfang W, Tyree MT, Steudle E (1999) Direct measurement of xylem pressure in leaves of intact maize plants. A test of the cohesion-tension theory taking hydraulic architecture into consideration. *Plant Physiology* 121(4):1191–1205.
27. Scoffoni C, et al. (2017) Outside-xylem vulnerability, not Xylem embolism, controls leaf hydraulic decline during dehydration. *Plant Physiology* 173(2):1197–1210.
28. Trifiló P, Raimondo F, Savi T, Lo Gullo MA, Nardini A (2016) The contribution of vascular and extra-vascular water pathways to drought-induced decline of leaf hydraulic conductance. *Journal of Experimental Botany* 67(17):5029–5039.
29. Zhu S (2020) Ph.D. thesis, Development of sensing framework for the soil-plant atmosphere continuum (Cornell Univeristy, Ithaca, NY, USA).
30. Hammer GL, et al. (2009) Can changes in canopy and/or root system architecture explain historical maize yield trends in the U.S. corn belt? *Crop Science* 49(1):299–312.
31. Zhuang J, Nakayama K, Yu GR, Urushisaki T (2001) Estimation of root water uptake of maize: An ecophysiological perspective. *Field Crops Research* 69(3):201–213.
